# Quantitative prediction of variant effects on alternative splicing using endogenous pre-messenger RNA structure probing

**DOI:** 10.1101/2021.09.13.460117

**Authors:** Jayashree Kumar, Lela Lackey, Justin M. Waldern, Abhishek Dey, David H. Mathews, Alain Laederach

## Abstract

Splicing is a highly regulated process that depends on numerous factors. It is particularly challenging to quantitatively predict how a mutation will affect precursor messenger RNA (mRNA) structure and the subsequent functional consequences. Here we use a novel Mutational Profiling (-MaP) methodology to obtain highly reproducible endogenous precursor and mature mRNA structural probing data in vivo. We use these data to estimate Boltzmann suboptimal ensembles, and predict the structural consequences of mutations on precursor mRNA structure. Together with a structural analysis of recent cryo-EM spliceosome structures at different stages of the splicing cycle, we determined that the footprint of the B^act^ complex on precursor mRNA is best able to predict splicing outcomes for exon 10 inclusion of the alternatively spliced *MAPT* gene. However, structure alone only achieves 74% accuracy. We therefore developed a β-regression weighting framework that incorporates splice site strength, structure and exonic/intronic splicing regulatory elements which together achieves 90% accuracy for 47 known and six newly discovered splice-altering variants. This combined experimental/computational framework represents a path forward for accurate prediction of splicing related disease-causing variants.

## Introduction

Precursor messenger RNA (pre-mRNA) splicing is a highly regulated process in eukaryotic cells (Z. Wang and Burge 2008). Numerous factors control splicing including *trans*-acting RNA-binding proteins (RBPs), components of the spliceosome, and the pre-mRNA itself. Pre-mRNA structure is a key attribute that directs splicing, particularly alternative splicing, but we have only a poor understanding of pre-mRNA structure-mediated splicing mechanisms (Taylor and Sobczak 2020). In addition, it is particularly challenging to develop quantitative models capable of predicting splicing outcome, specifically the Percent Spliced In (PSI) for alternatively spliced junctions. This difficulty is especially true for predicting the effects of genetic variation at exon-intron junctions. Indeed, mutations may affect not only the binding specificity of RBPs but also may alter pre-mRNA structure (Tazi, Bakkour, and Stamm 2009).

Similar to the challenge of predicting PSI outcomes, the consequences of mutations on pre-mRNA structure are difficult to predict. First and foremost, little is known about native pre-mRNA structure because pre-mRNAs are relatively short-lived in cells (Herzel et al. 2017). Only recently has pre-mRNA structure determination become amenable to high resolution in vivo experimental characterization (Mustoe et al. 2018). Second, it is not clear what structures of a pre-mRNA control spliceosome assembly and activity. Finally, we lack quantitative measures for the relative weighting of RBPs’ affinity for specific motifs in pre-mRNA to the importance of pre-mRNA structure. Several technical developments address these issues and enable us to propose an integrated, RNA structure based-framework that accurately predicts the percent of splicing. In this study, we used a combination of endogenous pre-mRNA chemical structure probing (Homan et al. 2014), an RNA structure model that considers multiple alternative structures in equilibrium (Dethoff et al. 2012; Lai et al. 2018), quantitative analysis of exonic and intronic splicing enhancers/silencers (Fairbrother et al. 2002; Z. Wang et al. 2004; Yang Wang, Ma, et al. 2012; Yang Wang, Xiao, et al. 2012), and a β-regression weighting (Ferrari and Cribari-Neto 2004).

In this work we measure endogenous pre-mRNA structure in vivo by combining recent developments in RNA structure Mutational Profiling (so-called -MaP approaches) with targeted amplification of specific exon-intron junctions. This novel approach enables us to obtain single-nucleotide RNA structure probing data for endogenous pre- and mature mRNAs in the same cell. The high reproducibility of these data also makes it possible to use Boltzmann suboptimal sampling guided by the data (Spasic et al. 2018) to predict free energies of unfolding for an ensemble of structures. In addition, we can now leverage recent high resolution cryo-Electron Microscopy (cryo-EM) structures of various stages of the spliceosome during the splicing cycle to reveal the effective spliceosomal footprint on pre-mRNA (L. Zhang et al. 2019).

As a model system to validate our framework, we study the effects of 47 experimentally measured mutations at the Exon10-Intron10 junction of the human Microtubule-Associated Protein Tau gene, *MAPT* (Park, Ahn, and Gallo 2016; Catarina Silva and Haggarty 2020). Exon 10 is a cassette exon that is alternatively spliced resulting in a Tau protein with either four microtubule binding repeats (4R) or three repeats (3R). The ratio of 3R to 4R isoforms is approximately 1:1 (Hefti et al. 2018). This is highly unusual for a splicing event as single-cell RNA-seq analysis demonstrates that this type of event, where alternative isoforms are expressed equally, comprises less than 20% of all splicing events (Song et al. 2017). The Exon10-Intron10 junction has 29 clinically validated disease-causing mutations (Stenson et al. 2003) that impair the function of Tau protein and are implicated in many neurodegenerative diseases (Spillantini et al. 1998; Hutton et al. 1998; Clark et al. 1998; Rizzu et al. 1999; Goedert et al. 1999). Although some mutations alter the Tau protein sequence (Mirra et al. 1999; Iseki et al. 2001), 20 disease-associated mutations are known that deregulate *MAPT* pre-mRNA splicing by altering the 1:1 ratio of 3R to 4R *MAPT* isoforms (Hutton et al. 1998; D’Souza et al. 1999; Hasegawa et al. 1999; Jiang et al. 2000). An additional 27 mutations were previously experimentally tested to measure Exon 10 PSI with splicing assays (D’Souza and Schellenberg 2000; Tan et al. 2019; Grover et al. 1999), making this junction the most experimentally characterized junction of clinical importance in the human genome and an excellent system for developing forward predictive models of splicing. Our work thus provides a framework for integrating endogenous pre-mRNA structure probing data with our current structural understanding of spliceosome assembly and *trans*-acting RBPs to achieve unprecedented quantitative prediction accuracy of the effect of mutations at structured exon-intron junctions.

## Results

### Median ratio of individual and tissue 3R to 4R MAPT mRNA isoforms is 1:1

Splicing of *MAPT* Exon 10 yields a 1:1 ratio of alternatively spliced isoforms (Goedert et al. 1989; Andreadis 2005). To corroborate the 1:1 isoform ratio among tissues and individuals, we analyzed RNA-sequencing data from the Genotype-Tissue Expression (GTEx) database (Lonsdale et al. 2013). We selected tissue types with median *MAPT* transcripts per million greater than 10 (Figure 1-figure supplement 1A) and calculated the Percent Spliced In (PSI) value for Exon 10 for each sample (Figure 1A-source data 1; Materials and methods). We examined the distribution of PSIs for each tissue type over 2,315 tissue samples in 375 individuals of median age 61 (Figure 1A; Figure 1-figure supplement 1B). A PSI of 0 indicated that none of the *MAPT* transcripts in a sample had Exon 10 spliced in (3R isoform), whereas a PSI of 1 corresponded to all *MAPT* transcripts having Exon 10 spliced in (4R isoform). We found variation in Exon 10 PSI both within and between different tissue types; the pituitary gland had the largest variation among brain tissues, and the cerebellum had the least variation but the difference between the two standard deviations was 0.04. Also, while the pituitary gland and caudate had the lowest and highest median Exon 10 PSI respectively among individual samples, the distance between the two values was only 0.25. Interestingly, although *MAPT*’s function in breast tissue is not understood compared with its function in the brain, for breast tissue, individuals had greater variation in Exon 10 PSI and a lower median PSI compared with the pituitary gland (Figure 1-figure supplement 1B). We also discovered a large amount of variation within tissues of an individual (Figure 1-figure supplement 1C), although there was significantly greater variation between individuals than within a single individual (see Supplementary file 1 for ANOVA table). Overall, 75% of samples were within a standard deviation of the median PSI of 0.54, which confirmed that the 3R to 4R isoform ratio was approximately 1:1 among individuals and within different tissue types. The consistency of this isoform ratio, despite the likely presence of different levels of RBPs, suggest that inherent sequence and structural features regulate splicing at this exon-intron junction. RNA structure regulates alternative splicing around exon-intron junctions (Warf and Berglund 2010; Buratti and Baralle 2004) and a hairpin structure at the exon 10-intron 10 junction is implicated in establishing the 3R to 4R 1:1 isoform ratio (Hutton et al. 1998; Varani et al. 1999; Grover et al. 1999; Donahue et al. 2006). Hence, we next used high-throughput chemical mapping techniques to interrogate the endogenous in vivo structure of the *MAPT* junction.

**Figure 1:**
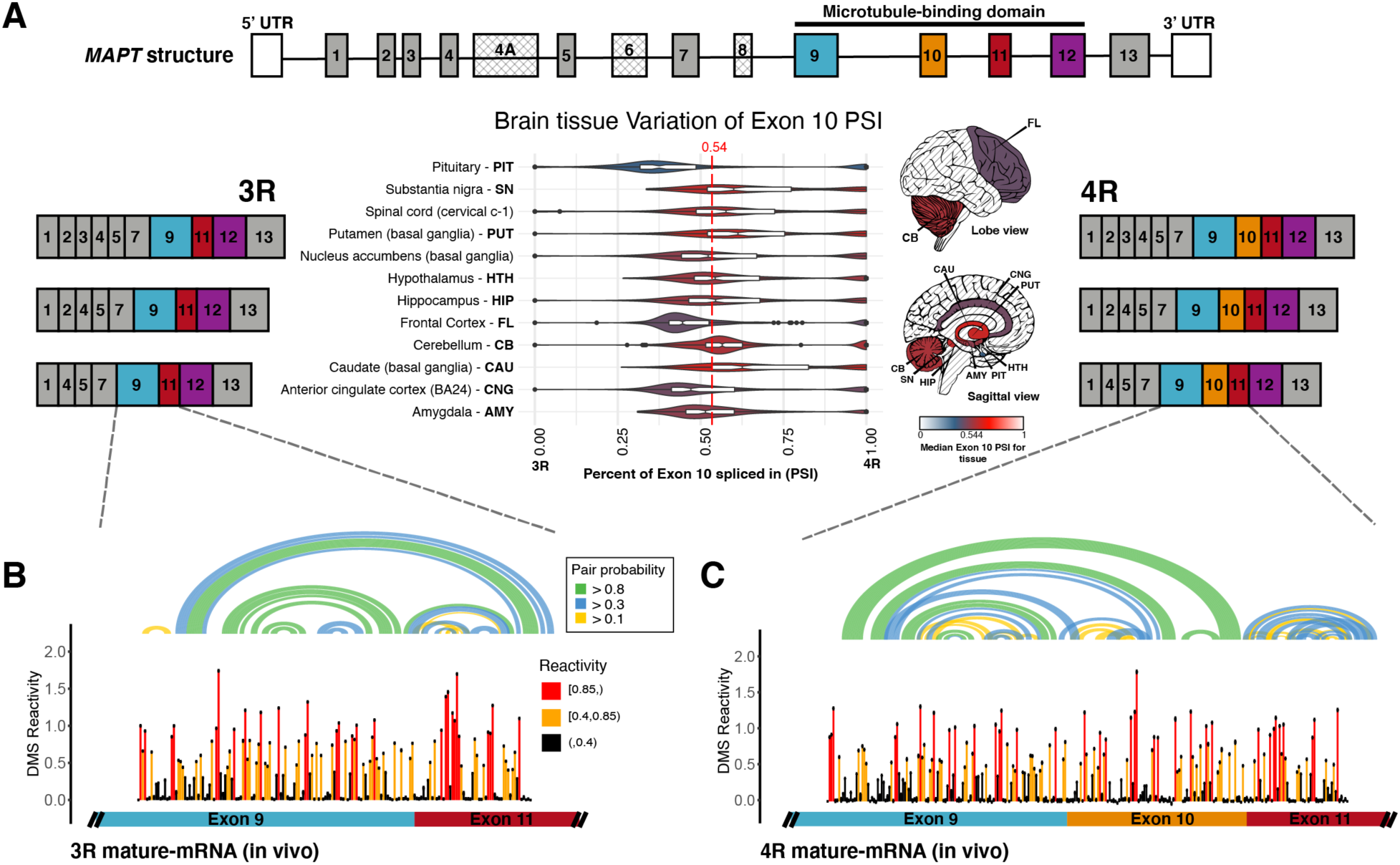
In vivo DMS-MaP structure probing data for 3R and 4R mature *MAPT* transcripts that are expressed in a 1:1 ratio. A) Ratio of 3R and 4R MAPT transcripts is approximately 1:1 among brain tissues. There are 14 exons alternatively spliced in *MAPT*. Exons 4A, 6, and 8 are not found in brain mRNA. The four exons highlighted in color are repeat regions that form the microtubule binding domain in the Tau protein. Exon 10 is alternatively spliced to form the 3 repeat (3R) or 4 repeat (4R) isoform. The six canonical transcripts found in the central nervous system can be separated into 3R and 4R transcripts. Percent Spliced In (PSI) of Exon 10 was calculated from RNA-seq data for 2315 tissue samples representing 12 central nervous system tissue types and collected from 375 individuals in GTEx v8 database. The violin plot for each tissue type and the corresponding region on the brain diagram is colored by the median PSI for all samples of a given type. The patterned regions on the brain diagram indicate tissue types with no data. Tissue types Spinal cord and Nucleus accumbens are not visualized on the brain diagram. The median PSI of 0.54 among all tissue samples is indicated by the red dotted line. B) In vivo DMS-MaP structure probing data across exon9-exon11 junction of 3R mature MAPT transcript. T47D cells were treated with DMS. Structure probing data for junctions of interest were obtained using primers (Supplementary file 4) following RT of extracted RNA. DMS reactivity is plotted for each nucleotide across spliced junctions. Each value is shown with its standard error and colored by reactivity based on color scale. High DMS reactivities correspond to unstructured regions, whereas low DMS reactivities correspond to structured regions. The base pairs of the predicted secondary structure guided by DMS reactivities are shown in the arcs colored by pairing probabilities. C) In vivo DMS-MaP structure probing data across exon9-exon10-exon11 junction of 4R mature MAPT transcript

### Structure of 3R and 4R MAPT mature mRNA isoforms is open and accessibility of exons is similar for the two isoforms

Although the structure of the *MAPT* pre-mRNA was previously studied computationally and in vitro (Varani et al. 1999; Lisowiec et al. 2015; Tan et al. 2019; Chen et al. 2019), the structures of the mature 3R and 4R isoforms and *MAPT* pre-mRNA have not been assessed in their endogenous in vivo context. We used dimethyl sulfate (DMS) to chemically probe RNA structure in T47D and neuronal SH-SY5Y cells and primer-amplified the Exon 9-Exon 11 and Exon 9-Exon 10-Exon 11 junctions during library preparation for Mutational-Profiling (-MaP) (Figure 1B; Materials and methods). This approach leverages the read-through aspect of MaP technology to probe the structure of two alternatively spliced isoforms in the same cells. DMS reactivities for replicates, and between T47D and SH-SY5Y MAPT mRNAs were highly correlated (Figure 1-figure supplement 2A; Figure 1-figure supplement 2B; Figure 1-figure supplement 2D; Figure 1-figure supplement 2E).

We also collected in vivo DMS data for the small ribosomal RNA (SSU) whose secondary structure is known from X-ray crystallography (Petrov et al. 2014) (Figure 1-figure supplement 3A). The DMS reactivities for unpaired nucleotides in the SSU were significantly higher than for paired nucleotides (Figure 1-figure supplement 3B), confirming our probing strategy accurately recapitulates RNA secondary structure. We used the SSU in vivo data to calibrate the estimation of equilibrium ensembles as guided by MaP technology (Methods and materials), and we validated that structure prediction guided by experimental DMS reactivities yielded more accurate estimation of the SSU structure (Figure 1-figure supplement 3C). The median DMS reactivity of the mature *MAPT* isoforms was 0.22, significantly greater than the median DMS reactivity of the SSU, 0.0083 (Figure 1-figure supplement 3D); these results suggested that the nucleotides of the mature *MAPT* isoforms were more accessible and unpaired compared with the highly structured SSU, indicating that our endogenous in vivo probing strategy reveals important differences in the structure of cellular RNAs.

Reactivities of Exon 9 and Exon 11 were highly correlated between the 3R and 4R isoforms (Figure 1-figure supplement 2C). Additionally, computed base-pairing probabilities guided by the experimental data for the two isoforms revealed that, although there were some long-range interactions, 66% of base pairs spanned less than 50 nucleotides and were contained within the exon units (Figure 1B). This result suggested that the mature exons function as their own structural unit. However, the mature isoform structures did not suggest how they might regulate splicing of Exon 10. Hence, we next chemically probed the MAPT pre-mRNA.

### MAPT pre-mRNA Exon 10-Intron 10 junction is more structured compared with the mature isoforms

Existence of a hairpin at the *MAPT* Exon 10-Intron 10 junction, implicated in regulating Exon 10 splicing, was established by NMR and in vitro chemical probing (Varani et al. 1999; Lisowiec et al. 2015); however, the endogenous in vivo structure of this region has yet to be determined. While collecting data for mature *MAPT* isoform junctions, we simultaneously obtained data for the pre-mRNA Exon 10-Intron 10 junction (Figure 2A; Materials and methods). Replicates were highly correlated (Figure 2-figure supplement 1A). Surprisingly, although Exon 10 was still being spliced, the reactivities for Exon 10 in pre-mRNA and the mature 4R isoform were highly correlated (Figure 2-figure supplement 1B). Again, base pairing between nucleotides appeared to be contained within exons, independent of introns. The reactivities were highly correlated between data collected in SH-SY5Y and T47D cells (Figure 2-figure supplement 1C); thus, despite likely differences in RBP concentrations, the structure of the pre-spliced region is the same between cell lines. Additionally, we found lower DMS reactivities for the pre-mRNA Exon 10-Intron 10 junction compared with the mature isoform junctions (Figure 2- figure supplement 1D), which suggests that pre-mRNA is more structured than mature mRNA. We uncovered strong evidence for the previously in vitro identified hairpin structure in the DMS reactivity data; pairing probabilities were greater than 0.8 for the entire hairpin stem (Figure 2A).

**Figure 2:**
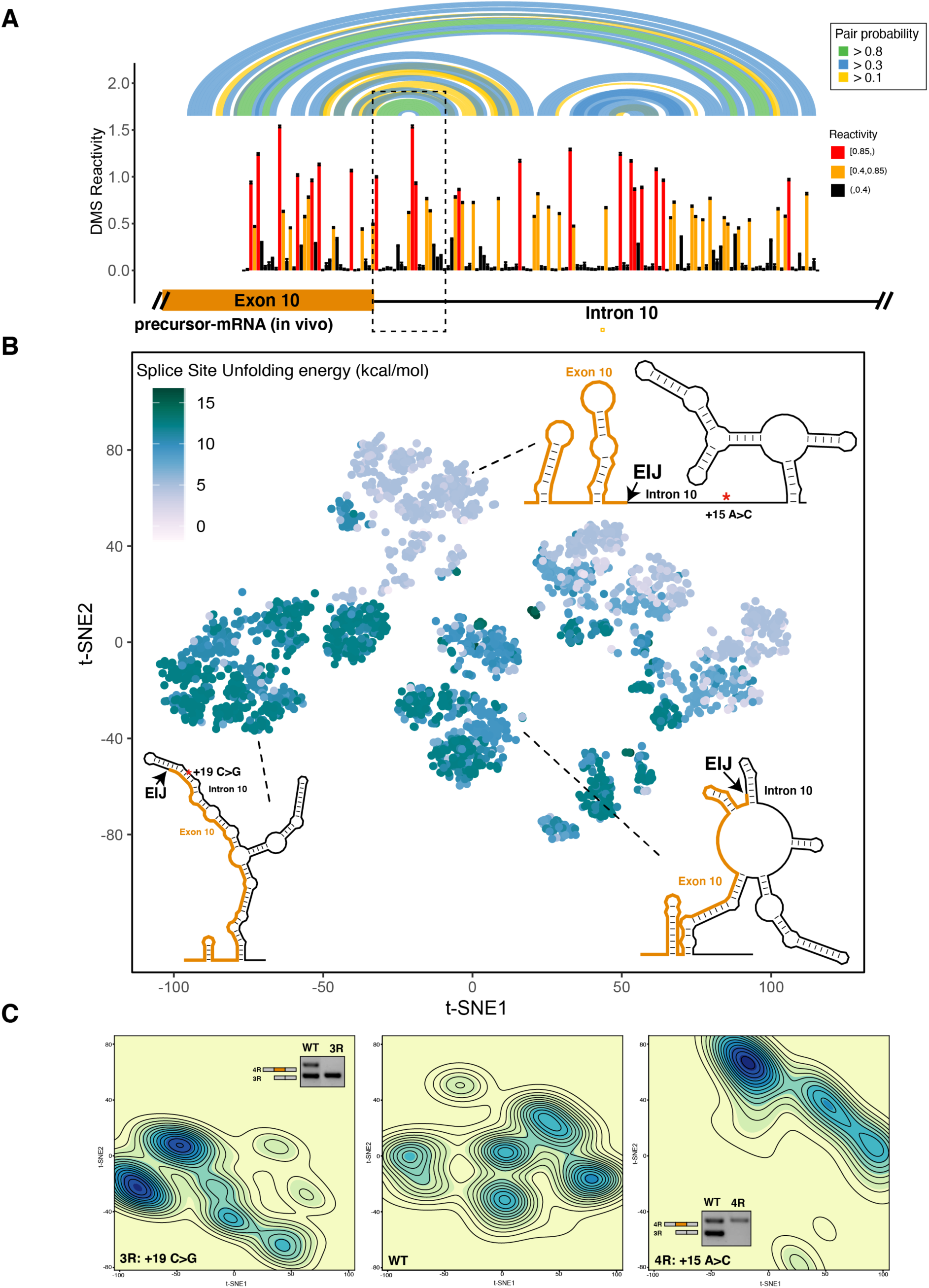
The 4R and 3R mutations shift DMS reactivity-guided structural ensemble of Exon 10-Intron 10 junction to more open and closed structures, respectively. A) In vivo DMS-MaP structure probing data across Exon 10-Intron 10 junction of precursor MAPT transcript. T47D cells were treated with DMS. Structure probing data for junctions of interest were obtained using primers (Supplementary file 4) following RT of extracted RNA. DMS reactivity is plotted for each nucleotide. Each value is shown with its standard error and colored by reactivity based on the color scale. High DMS reactivities correspond to unstructured regions, whereas low DMS reactivities correspond to structured regions. Base pairs of predicted secondary structure guided by DMS reactivities are shown by arcs colored by pairing probabilities. Strongly predicted hairpin structure near exon-intron junction is highlighted by dotted box. B) t-SNE Visualization of structural ensemble of wildtype (WT) and, +19C>G (3R) and +15A>C (4R) mutations. Structures were predicted using Boltzmann suboptimal sampling and guided by DMS reactivity data generated in A. Data were visualized using t-Distributed Stochastic Neighbor Embedding (t-SNE). Shown are 3000 structures corresponding to 1000 structures per sequence. Each dot represents a single structure and is colored by calculated unfolding free energy of splice site at exon-intron junction (3 exonic, 6 intronic bases). Data were clustered by k-means clustering and representative structures for three of the clusters are shown. The exon-intron junction is marked by EIJ on each structure. Positions of 3R and 4R mutations are marked by a red asterisk on their respective representative structures. C) Density contour plots of structural ensemble of WT and, 3R and 4R mutations. Contour plots were derived from the distribution of points on the t-SNE plot in B. The darker the blue, the higher the density of structures. Contour lines additionally represent density of points. Color scales for the three plots are identical. Gel insets of RT-PCR products from splicing assays in HEK293 cells for 3R and 4R mutation are in their respective density plots.

### Shifts in structural ensemble of MAPT Exon 10-Intron 10 junction associated with disease mutations correlate with changes in splicing level of Exon 10

Many RNAs inhabit multiple conformations in vivo to form a structural ensemble instead of a single rigid structure (Halvorsen et al. 2010; Adivarahan et al. 2018). We posit that a structural ensemble at the *MAPT* Exon 10-Intron 10 junction regulates Exon 10 splicing and disease mutations alter the composition of the structural ensemble to disrupt splicing.

We used Boltzmann sampling of RNA structures guided by DMS reactivity data (Spasic et al. 2018) (Materials and methods) to sample 1000 structures for the wild type and two mutant intronic sequences, +15A>C and +19C>G. The two mutations alter, in opposite directions, the isoform ratio at this junction (Tan et al. 2019). We visualized the structural ensemble for the 3000 structures using t-Distributed stochastic neighbor embedding (t-SNE) (Van Der Maaten and Hinton 2008) (Figure 2B; Materials and methods). Each structure is a dot and is colored by the ΔG^‡^ of unfolding of the 5’ splice site defined as the last three nucleotides of Exon 10 and the first six nucleotides of Intron 10 (Yeo and Burge 2004). The lower the unfolding free energy, the easier to unfold the structure. Overall, although there was a range of unfolding free energies for the three ensembles, there were three distinct populations of free energies for the three sequences (Figure 2-figure supplement 2A). We used k-means clustering to identify representative structures for each cluster (Figure 2B; Figure 2-figure supplement 2B; Materials and methods). We quantified and visualized the density of the clusters (Figure 2C; Materials and methods) and revealed distinct regions in the structure space occupied by each sequence. More than 55% of structures in the ensemble of the +19C>G mutation, which shifts the isoform balance entirely 3R (3R mutation) (Figure 2C inset), clustered in the lower left quadrant with larger unfolding free energies for the splice site. This result was evidenced by the highly base-paired exon-intron junction in the representative structure for the cluster. Hence, in the presence of the 3R mutation, the structural ensemble of the junction shifted towards more closed structures.

Conversely, greater than 50% of structures in the ensemble of the +15 A>C mutation, which shifts the isoform balance entirely 4R (4R mutation) (Figure 2C inset), were clustered in the upper left quadrant with lower unfolding free energies for the splice site. The representative structure for this region was more open and accessible around the exon-intron junction. Correspondingly, the wild-type sequence had structures distributed across the entire space consistent with an ensemble of structures. The exon-intron junction of the representative structure for this region was not as accessible with the 4R mutation, but it had fewer base-pairs than with the 3R mutation, a result recapitulated by the two other representative structures in the right quadrants (Figure 2-figure supplement 2B).

### Unfolding mRNA within the spliceosome B^act^ complex yields best prediction of Exon 10 splicing level

RNA structure controls alternative splicing by hindering or aiding accessibility of key regulatory regions to spliceosome components (McManus and Graveley 2011; Warf and Berglund 2010). The 5’ splice site, defined as the last 3 nucleotides of the exon and first 6 nucleotides of the intron, is the minimum region of RNA that must be accessible for base pairing with the U1snRNA (Blanchette and Chabot 1997; Singh, Singh, and Androphy 2007). However, the splicing cycle, orchestrated by the spliceosome, traverses multiple stages to prepare the pre-mRNA and catalyze the two-step splicing reaction (Matera and Wang 2014) (Figure 3A). The RNA itself adopts many conformations as different components of the spliceosome bind to it (L. Zhang et al. 2019). In addition to the 5’ splice site, a larger segment of RNA likely needs to unpair to accommodate the changing conformations induced by the spliceosome. We analyzed high resolution Cryo-EM structures of the human spliceosome Pre-B, B, Pre-B^act^, and B^act^ complexes (Charenton, Wilkinson, and Nagai 2019; Bertram et al. 2017; Townsend et al. 2020; X. Zhang et al. 2018) to quantify the number of nucleotides around the 5’ splice site for which sufficient density was observed in the cryo-EM structure and which were unpaired (Materials and methods). As can be seen in Figure 3A, the number of unpaired pre-mRNA nucleotides observed in each structure increased through the splicing cycle. Thus, it is likely that RNA structures outside the U1snRNA binding site have to be unfolded to accommodate splicing.

**Figure 3:**
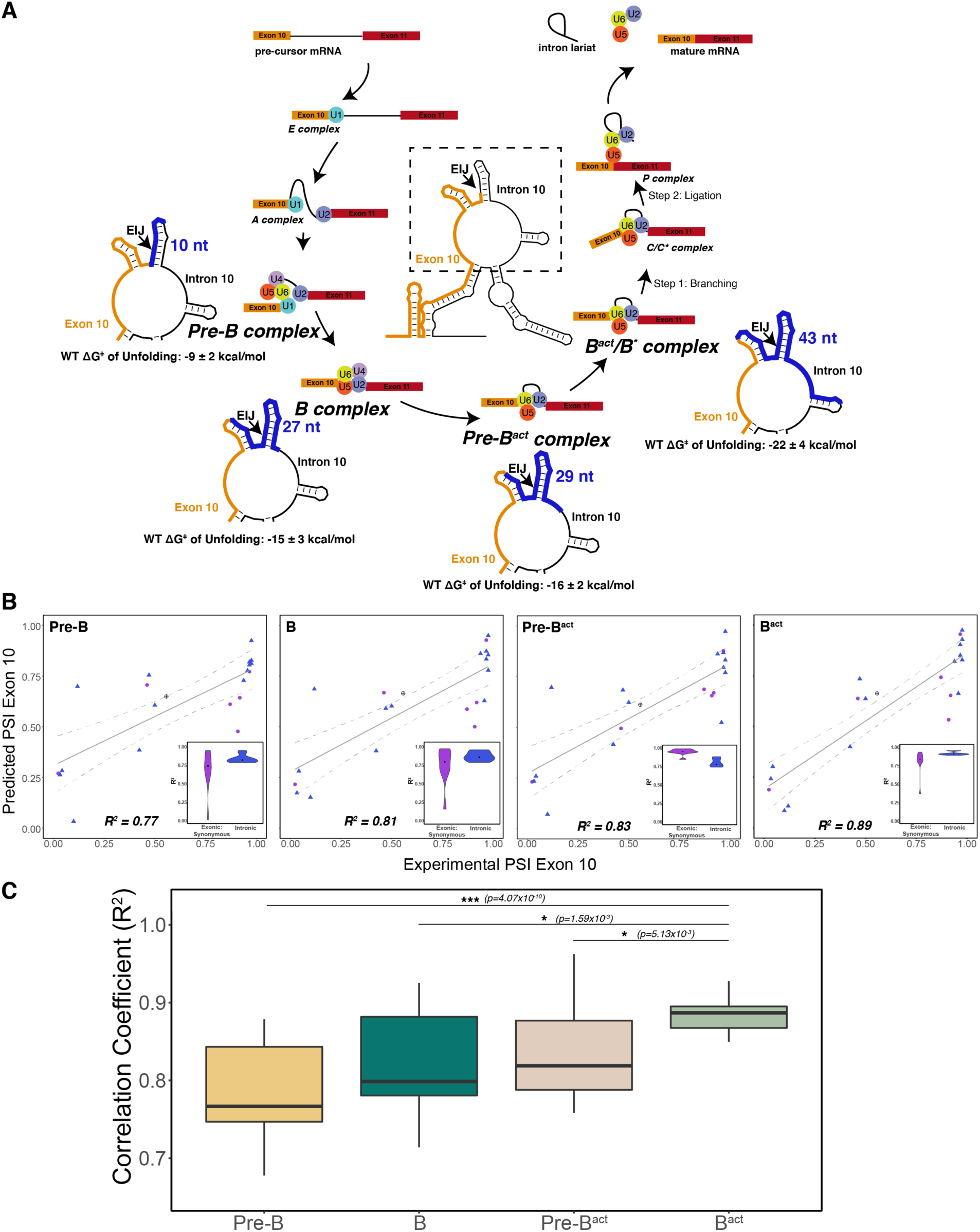
The best predictor of Exon 10 PSI for intronic and synonymous mutations was the unfolding free energy of pre-mRNA during the B^act^ stage of splicing. A) Spliceosome footprint on pre-mRNA during splicing cycle. Structure in the center of the cycle is the WT representative structure from Fig 2B. The dotted box indicates the zoomed-in region at each stage of interest. Cryo-EM structures of the human spliceosome complex at stages Pre-B (PDB ID: 6QX9), B (PDB ID: 5O9Z), Pre-B^act^ (PDB ID: 7ABF) and B^act^ (PDB ID: 5Z56) are available in the Protein Data Bank. The region around the 5’ splice site of pre-mRNA within the spliceosome at each stage is highlighted in blue on the zoomed-in representative structure. The number of nucleotides for each stage is as follows: Pre-B (2 exonic, 8 intronic); B (10 exonic, 17 intronic); Pre-B^act^ (9 exonic, 20 intronic); B^act^ (12 exonic, 31 intronic). These values represent the minimum number of nucleotides required to be unfolded to be accessible to the spliceosome. The mean free energy and standard error to unfold RNA within the spliceosome at each stage is calculated for the WT structural ensemble and indicated under the zoomed-in structure. B) Exon 10 PSIs of synonymous and intronic mutations predicted with the unfolding free energy of pre-mRNA within the spliceosome in B, Pre-B, Pre-B^act^, B^act^ stages versus corresponding experimental PSIs measured in splicing assays. Exon 10 PSIs were predicted using Eq. 1. Grey line represents the best fit with dotted lines indicating the 95% confidence interval. Pearson correlation coefficients (R^2^) of experimental to predicted PSIs were calculated for each stage. Violin plots (inset) show R^2^s calculated for each mutation category by training and testing on subsets of all mutations by non-parametric bootstrapping; Synonymous (n=6), Intronic (n=14), Wildtype (n=1). C) Overall Pearson correlation coefficients (R^2^) calculated for experimental versus predicted Exon 10 PSIs by nonparametric bootstrapping of mutations. Subsets of mutations were randomly sampled 10 times, trained and tested using unfolding free energy of the exon-intron junction region of pre-mRNA within the spliceosome for the respective splicing stage. Pearson’s R^2^ was calculated by comparing predicted PSIs to experimental PSIs. A two-tailed Wilcoxon Rank Sum test was used to determine significance between B^act^ complex and the other three complexes. Level of significance: ***p-value < 10^-6^, **p-value < 0.001, * p-value < 0.01

To evaluate the footprint of the spliceosome that best predicts splicing outcome, we initially focus on predicting 20 synonymous and intronic mutations as a training set (Figure 3-figure supplement 1A). These mutations are most likely to have a structural component to their function (Sharma et al. 2019; Lin, Taggart, and Fairbrother 2016). The distribution of ΔG^‡^ of unfolding of the splice sites in the presence of these mutations was correlated with Exon 10 PSI (Figure 3-figure supplement 1B). We calculated the ΔG^‡^ of unfolding of the RNA near the 5’ splice site in the four splicing stages’ footprints. Features of the unfolding free energy distribution including mean and standard deviation were then used in a beta regression to predict Exon 10 PSI (Materials and methods; Eq. **1**). Unfolding larger regions of the exon-intron mRNA junction improved the predictive power of the model, and the B^act^ complex footprint yielded the best prediction accuracy (R^2^ = 89%; Figure 3B). Crucially, we found that using features of the distribution of unfolding free energies in the structural ensemble produced greater predictive power than simply using the unfolding free energy of a single minimum free energy (MFE) structure (Figure 3-figure supplement 1C). We performed bootstrapping cross-validation and confirmed that unfolding the RNA within the B^act^ spliceosome complex yielded the best prediction (Figure 3C). We tested the structural ensemble-based model on 24 non-synonymous and compensatory mutations. Although the model performed well for compensatory mutations (median bootstrapped R^2^=0.76), it yielded significantly less accurate predictions for non-synonymous mutations (median bootstrapped R^2^=-0.21) (Figure 3-figure supplement 1D). One possible reason this structure-only model has limited performance is that it does not account for the effects of mutations on potential splicing regulatory elements (SREs) in the sequence.

### Effect of exonic non-synonymous mutations was best predicted by motif strength changes of splicing regulatory elements

Exon 10 splicing is highly regulated by differential binding of RBPs to *cis*-SREs within exon 10 and intron 10 (Qian and Liu 2014). While our structure-only model performs moderately well for 47 mutations (R^2^=0.74) (see Supplementary file 2 for further details about mutations), *MAPT* Exon 10 PSIs of non-synonymous mutations were poorly predicted (median bootstrapped R^2^ = -0.21, Figure 4-figure supplement 1B). Hence, we investigated whether these non-synonymous mutations are predicted better by incorporating changes to the strength of adjacent SREs. We identified SREs by similarity to reported general enhancer and silencer hexamer motifs (Fairbrother et al. 2002; Z. Wang et al. 2004; Yang Wang, Ma, et al. 2012; Yang Wang, Xiao, et al. 2012) (Materials and methods). We calculated the changes to splice site, enhancer, and silencer motif strengths in the presence of a mutation (Materials and methods) and visualized the motif strength changes in a heatmap (Figure 4A). We found that using splice site strength as the sole predictor yielded poor prediction of Exon 10 PSI in all mutation categories (Figure 4B; Eq. 3) because most mutations were outside the splice site. We quantified a weak positive correlation between PSI and enhancer strength and a significant negative correlation between PSI and silencer strength (Figure 4A; Figure 4-figure supplement 1C). We modeled Exon 10 PSI with the changes to the motif strength of all splicing regulatory elements (Eq. 4) and found an increase in prediction accuracy (R^2^=0.51; Figure 4C) compared with using only splice site strength (R^2^=0.29). Non-synonymous mutations were predicted more accurately using SRE strength with a median bootstrapped R^2^ of 0.75.

**Figure 4:**
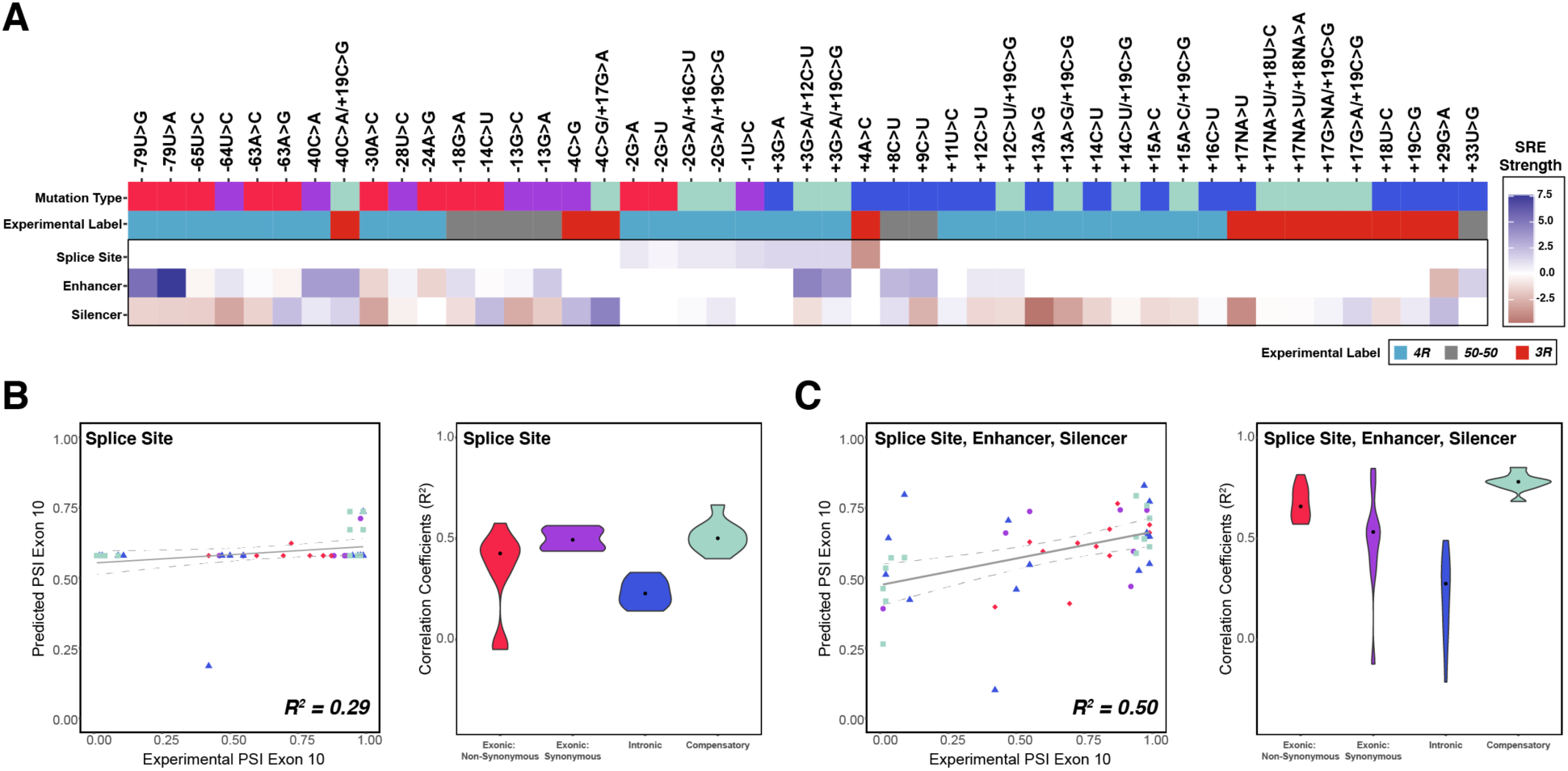
Combining the strength of all splicing regulatory elements improves prediction of Exon 10 PSI by 75% compared with using only splice site strength. A) Heatmap of splicing regulatory element (SRE) relative strength for 47 mutations compared with wildtype (WT). A value of 0 indicates mutation does not change WT SRE strength, positive values indicate SRE strength is greater than WT, and negative values indicate SRE strength is weaker than WT. Splice site strengths were calculated using MaxEntScan; a splice site was defined as the last 3 nucleotides of the exon and first 6 nucleotides of the intron. Enhancer and silencer strengths were calculated from position weight matrices of known motifs derived from cell-based screens (Materials and methods). B) Exon 10 PSIs of 47 mutations predicted from change in splice site strength and plotted against experimental PSIs measured in splicing assays. Exon 10 PSIs predicted using Eq. 3. Each point on the scatterplot represents a mutation and is colored by mutation category. Grey line represents the best fit with dotted lines indicating the 95% confidence interval. Pearson correlation coefficient (R^2^) calculated of experimental to predicted PSIs. Violin plot shows R^2^s calculated for each category by training and testing on subsets of all mutations by non-parametric bootstrapping; Exonic non-synonymous (n=11), Exonic synonymous (n=7), Intronic (n=15), Compensatory (n=14), Wildtype (n=1). C) Exon 10 PSIs of 47 mutations predicted by combining change in splice site, enhancer, and silencer strength and plotted against experimental PSIs measured in splicing assays. Exon 10 PSIs predicted using Eq. 4.

Many RBPs have been identified that regulate *MAPT* Exon 10 splicing (Qian et al. 2011; Ian D’Souza and Schellenberg 2006; Kondo et al. 2004; J. Wang et al. 2004; L. Gao et al. 2007; S. Ding et al. 2012; Broderick, Wang, and Andreadis 2004; Yan Wang et al. 2010; Kar et al. 2006, 2011; P. Ray et al. 2011). To determine whether these proteins specific to Exon 10 splicing would improve the model’s accuracy, we calculated changes to the strength of their RBP motifs obtained from high throughput sequencing of bound RNAs (Dominguez et al. 2018; D. Ray et al. 2013) (Materials and methods). Unlike SRE motifs, there was no clear pattern or correlation between motif strength change and PSI (Figure 4-figure supplement 2A, B). Subsequently, the model’s prediction accuracy was lower (R^2^=0.08, Figure 4-figure supplement 2C), and changes to the strength of general SRE motifs were better predictors of Exon 10 PSI.

### Model with both structural and SRE motif changes yields best prediction of Exon 10 PSI

Our quantitative models showed that, although SRE motif changes accurately predicted the effects of non-synonymous mutations, structural changes were a better predictor of splicing outcomes of intronic and synonymous mutations. Combining all features (Eq. 6) yielded the highest prediction accuracy (R^2^ = 0.89) (Figure 5A). This combined interactive model consistently produced more accurate predictions of Exon 10 PSI compared with a structure-only model and an SRE-only model for all mutation categories (Figure 5B). An additive model (Eq. 7) had lower prediction accuracy (R^2^= 0.80) (Figure 5-figure supplement 1A), and this lower accuracy resulted primarily from less accurate PSI predictions of non-synonymous mutation effects (Figure 5-figure supplement 1B).

**Figure 5:**
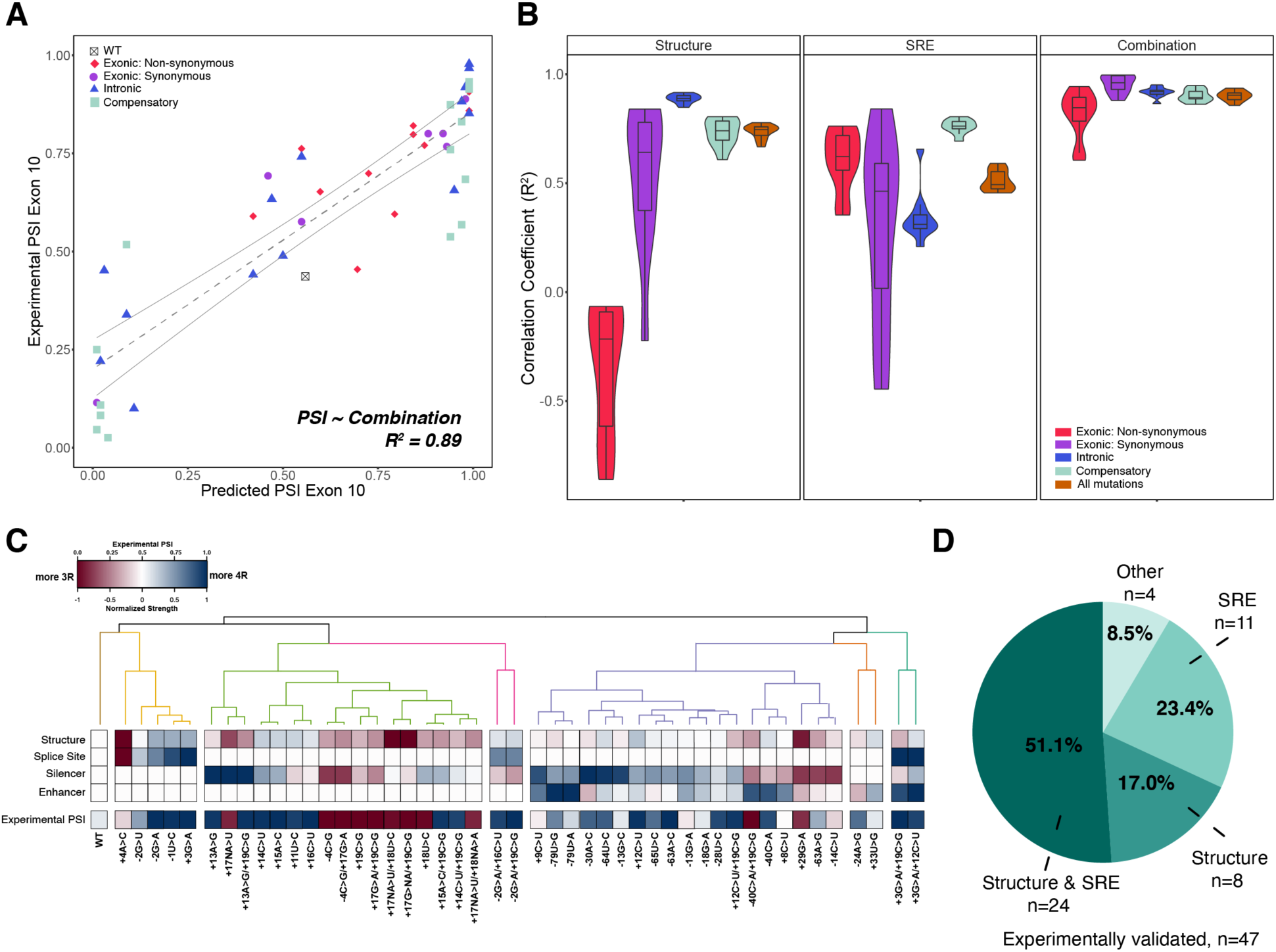
Combining structure and SRE strength into a unified model is the best predictor of Exon 10 PSI. A) Exon 10 PSIs of 47 mutations predicted from combined model using structure and SRE strength and fit to experimental PSIs measured in splicing assays. Exon 10 PSIs predicted using Eq. 6. Each point on scatterplot represents a mutation and is colored by mutation category. Grey line represents the best fit with dotted lines indicating the 95% confidence interval. Pearson correlation coefficient (R^2^) calculated of experimental to predicted PSIs. B) Violin plots of correlation coefficients for each mutation category for structure model, SRE model, and combined model. R^2^s calculated for each mutation category by training and testing on subsets of all mutations by non-parametric bootstrapping 10 times. Structure model uses unfolding free energy of pre-mRNA within spliceosome at B^act^ stage as predictor. SRE strength model uses relative change in SRE strength as predictor. Combined model using both structure and SRE strength and weighs the features based on if mutation is intronic/synonymous or non-synonymous (Materials and methods). C) Heatmap of the normalized changes in structure and SRE strength for each mutation clustered by affected features. Features were normalized such that a value of 1 implied that change in the feature should result in Exon 10 being spliced in (4R isoform, blue), whereas a value of 0 implies Exon 10 should be spliced out (3R isoform, red). Mutations were clustered using hierarchal clustering on normalized features (Materials and methods). Experimental PSIs are plotted for each mutation with a PSI of 1 colored as blue, PSI of 0.5 colored as white and PSI of 0 colored as red. D) Pie chart showing the features that regulate Exon 10 splicing for the 47 experimentally validated mutations. The pie chart was generated based on the heatmap in C. Exon 10 splicing for 51.1% of mutations is supported by changes in both structure and SRE, which implies that structure, at least one SRE, and PSI are either all blue or all red. Exon 10 splicing for 23.4% of mutations is supported by changes in SRE wherein one of the SREs is the same color as PSI. For 17.0% of mutations, structural changes support Exon 10 splicing wherein structure and PSI are the same color. For 4 mutations (8.5%), the colors of none of the features match the color of PSI.

To determine whether structure or SRE changes were responsible for the splicing changes from each mutation, we hierarchically clustered the four primary features for the 47 experimentally validated mutations (Materials and methods). Six categories emerged from the clustering of features (Figure 5C) where approximately 80% of mutations modified both structure and silencer strength (Figure 5-figure supplement 1C). Further, we found that for more than 50% of mutations both structure and SRE motif strength were altered in the same direction and accordingly promoted Exon 10 splicing in that direction (Figure 5D). For the remaining mutations in which structure and SRE strength changed in opposite directions, structure dominated the direction of splicing for 18% of mutations, and SRE strength was dominant for 20% (Figure 5D). Overall, these results supported our conclusion that both structure and SREs have equally important effects in regulating splicing at this exon-intron junction.

### Mutations around the MAPT Exon 10-Intron 10 junction skew to Exon 10 inclusion

Having established that our quantitative models accurately predicted Exon 10 PSIs for experimentally validated mutations, we interrogated the model by performing a systematic mutagenic analysis spanning a 100-nucleotide window of the exon-intron junction (Figure 6A). Our model predicts that more mutations result in the inclusion of Exon 10 (4R isoform). This is consistent with the observation that a majority (75%) of known disease associated mutations (Figure 6B) are also 4R; this result is also consistent when categorized by all substitution types (Figure 6-figure supplement 1A). We found that a significantly greater proportion of disease mutations (76.4%) resulted in changes to both structure and SRE compared with non-disease mutations (36.0%) (Figure 6C) suggesting that mutations which affect both structure and SREs have a greater likelihood of causing disease compared with mutations that alter only one of the two factors. Intriguingly, mutations overall caused a slight skew towards a structured exon-intron junction, which would result in decreased inclusion of Exon 10 (Figure 6A, Figure 6-figure supplement 1B). However, changes to SRE strength skewed towards increased inclusion of Exon 10 (Figure 6-figure supplement 1C), which suggested that SREs were acting to counter the effect of structural changes. Our model reveals how a complex balance of structure and SRE RBP binding sites effectively results in the observed 50:50 ratio of the 3R and 4R isoforms.

**Figure 6:**
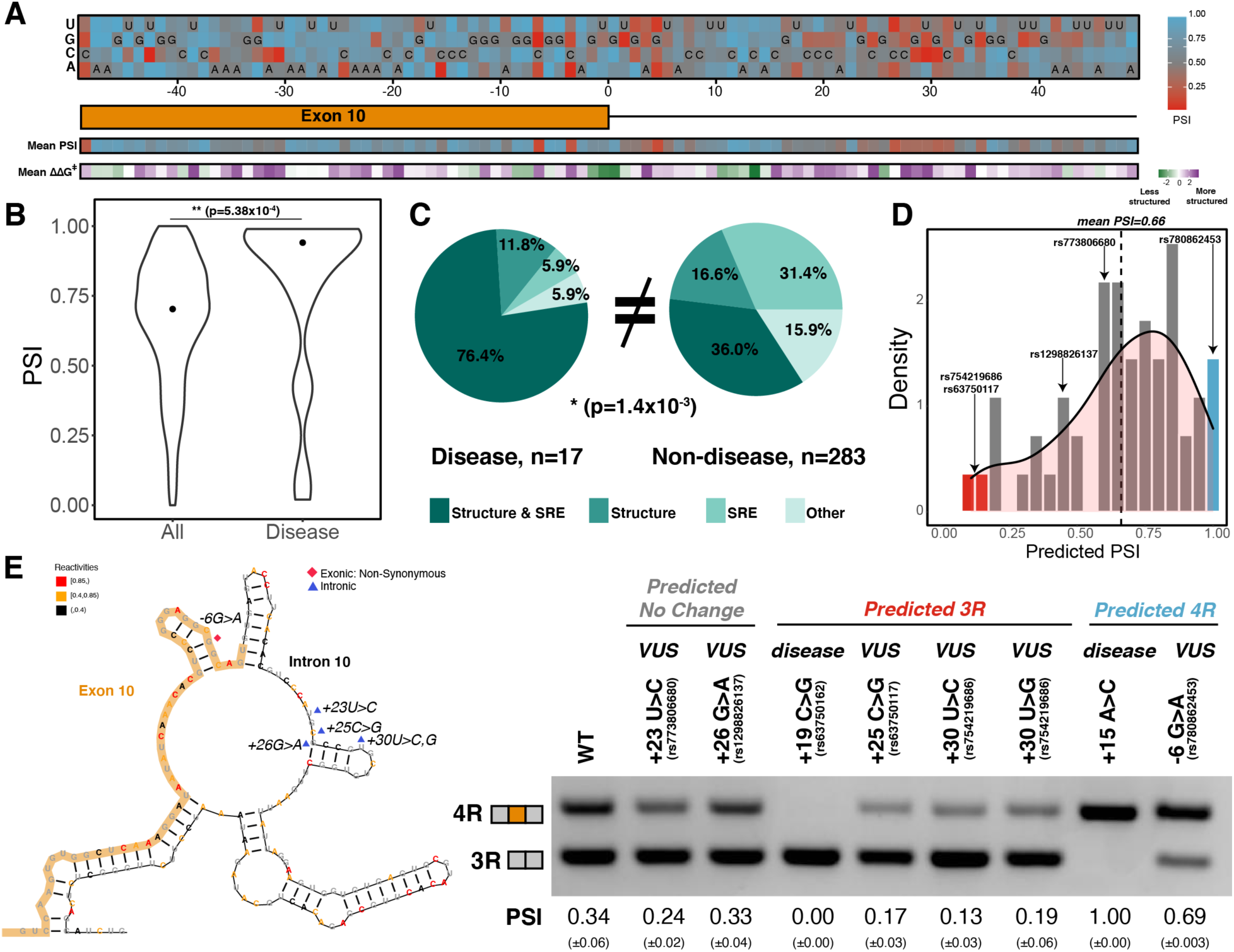
Mutations around Exon 10-Intron 10 junction skew towards inclusion of Exon 10. A) Heatmap of predicted Exon 10 PSIs for every possible mutation around 100 nucleotide window of Exon 10-Intron 10 junction. Combined model was trained using 47 mutations with experimental PSIs measured from splicing assays and used to predict PSIs for all mutation combinations for 100 nucleotides around the junction. Tiles with sequence indicate the wild type nucleotide at the position. Heatmap of mean PSI per position and mean relative change in unfolding free energy of pre-mRNA within spliceosome at B^act^ stage compared with wild type is shown below the gene diagram. B) Violin plot of predicted PSIs for all possible mutations around Exon 10-Intron 10 junction and only disease mutations. All possible mutations (n=300), disease mutations (n=17). A two-tailed Wilcoxon Rank Sum test was used to determine significance between the two categories. Level of significance: ***p-value < 10^-6^, **p-value < 0.001, * p-value < 0.01 C) Pie chart showing features that drive Exon 10 splicing for disease and non-disease mutations. The pie chart was generated by quantifying the number of mutations for which the direction of predicted Exon 10 PSI matched the direction of structure or SRE change. Exon 10 splicing for 76.4% of disease mutations is supported by changes to both structure and SRE compared with only 36.0% of non-disease mutations. The difference in proportions was tested with a one-tailed Fisher’s exact test. D) Histogram displaying the distribution of predicted PSIs using the combined model for 55 variants of unknown significance (VUSs) found in dbSNP. Density curve was overlaid on top of histogram showing that predicted PSIs skew away from 3R. Dotted line shows mean predicted PSI of 0.66. VUSs tested in splicing assays are indicated by their dbSNP RS IDs. E) Representative gel of RT-PCR data for splicing assay in the presence of VUSs. Splicing reporter was transfected into HEK293 cells. The mean Exon 10 PSI displayed for each variant was calculated from three replicates and standard error is shown in brackets below. Structure diagram on left displays the location of the VUSs tested.

To assess the general applicability of our model beyond our mutation training set, we predicted Exon 10 PSIs for 55 variants of unknown significance (VUSs) found in dbSNP (see Supplementary file 3 for further details of VUSs). These are mutations observed in the human population but are not currently associated with disease. The mean Exon 10 PSI for VUSs was 0.66, and 70% were within a standard deviation of the mean (Figure 6D). We observed that only a few mutations were predicted to have a PSI of zero (3R) (Figure 6D red bar). We therefore experimentally verified with splicing assays (Materials and methods) 6 VUSs: 3 VUSs predicted to be 3R, 1 VUS predicted to be 4R and 2 VUSs predicted to maintain the WT splicing ratio (Figure 6D). We found these 6 predictions were correct (Figure 6E). The three 3R VUSs made the region around the exon-intron junction more structured while the 4R VUS made the region less structured compared to WT (Figure 6-figure supplement 1D) matching the direction of Exon 10 splicing change. Though we see changes to SRE strength match up to Exon 10 splicing direction for +30U>C and -6G>A, this was not the case for +25C>G and +23U>C (Figure 6-figure supplement 1E). For +23U>C and +26G>A, we observed changes in structured-ness around the exon-intron junction and silencer strengths in diverging directions (Figure 6-figure supplement 1D, E) suggesting that these opposing changes would preserve the WT 3R/4R ratio.

## Discussion

### In vivo DMS chemical probing of endogenous MAPT Exon 10-Intron 10 junction

Splicing specificity is complex (Baralle and Giudice 2017). The spliceosome does not rely on sequence alone to correctly identify 5’ and 3’ splice sites; other cues ensure correct binding to appropriate locations. In addition, the 5’ splice site must be accessible to permit base pairing with the U1snRNA to initiate splicing (Roca et al. 2012). The *MAPT* Exon 10-Intron 10 junction is a well-studied example of the effect of 5’ splice site secondary structure in splicing regulation. A hairpin was hypothesized initially because disease mutations close to the exon-intron junction (Hutton et al. 1998; Grover et al. 1999) shifted the isoform balance to either completely exclude or include Exon 10. Although NMR, in vitro chemical probing, and computation confirmed the presence of the hairpin (Varani et al. 1999; Chen et al. 2019; Lisowiec et al. 2015), recent studies showed that most RNAs were less structured in vivo and in the nucleus compared with in vitro conditions (Sun et al. 2019; Rouskin et al. 2014). However, our results revealed that this is not the case for the Exon10-Intron10 junction: in vivo chemical probing of the endogenous junction showcased strong evidence of structure.

In this study we observed that, in vivo, endogenous exons are less structured than introns, as found by Sun et al (Sun et al. 2019). Mature *MAPT* 3R and 4R exon-exon junctions are less structured compared with the pre-mRNA Exon 10-Intron 10 junction. The high correlation of structure we observed between the same exons found in different *MAPT* isoforms corroborates results observed with yeast ribosomal protein genes (Zubradt et al. 2016), which suggests that RNA folding in both pre- and post-spliced human exons is highly local and modular in exons.

### Changes to structural ensemble around the 5’ splice site are strong predictors of Exon 10 splicing

We showed that structural ensembles have an important function at the Exon 10-Intron 10 junction. If the 5’ SS was always paired, only one isoform lacking Exon 10 would result. However, the simultaneous presence of 3R and 4R isoforms implies that the junction is accessible in a subset of the structures. Unlike transfer RNAs and ribosomal RNAs that have single structures (Petrov et al. 2014), most RNAs are dynamic, unfolding and refolding within a landscape (Cruz and Westhof 2009; Giegé et al. 2012). We found disease mutations produced distinct shifts in the ensemble of the *MAPT* Exon 10-Intron 10 junction; the shifts corresponded to changes in the 3R:4R isoform ratio and confirmed that ensembles are essential at this junction. The activity of ensembles was corroborated by our quantitative model; including free energy features of the structural ensemble produced 1.5 times more accurate prediction of Exon 10 PSI compared with the unfolding free energy of the minimum free energy (MFE) structure.

### Considering a larger spliceosome footprint on pre-mRNA produced more accurate prediction of Exon 10 PSI

The U1snRNA base pairs with the nine nucleotide sequence around the exon-intron junction (Roca et al. 2012). However, our analysis of the Cryo-EM structures of the human spliceosomal assembly cycle revealed that a larger region of the pre-mRNA interacts with the spliceosome during the splicing cycle and is therefore unfolded. Like the other main cellular ribonucleoprotein complex, the ribosome (Ingolia 2016), there is likely a spliceosomal footprint on the pre-mRNA and a minimum span around the splicing signals (5’ splice site, 3’ splice site and branch point) must be single-stranded for splicing to occur. Accordingly, our structural model performed most accurately when we used the unfolding free energy of 43 nucleotides around the 5’ exon-intron junction that exists within the spliceosome B^act^ complex. This suggests that structures distal to the exon-intron junction regulate Exon 10 splicing, a finding that corroborates evidence that RNA structure near this exon-intron junction is more extended than previously determined (Tan et al. 2019). This result, combined with our use of Boltzmann suboptimal sampling demonstrates the key role of pre-cursor mRNA structure in splicing outcome.

### RNA structure and SREs have complementary functions in MAPT Exon 10 regulation

Considerable evidence supports a function for either splicing regulatory elements and their corresponding RBPs or RNA structure in alternative splicing of *MAPT* Exon 10 at the 5’ splice site (Andreadis 2012). However, there was no consensus as to which of the two factors is dominant. The regression model we developed established the relative importance of RNA structure vs. SREs at the exon-intron junction. We discovered a cooperative relationship between SREs and RNA structure whereby exonic non-synonymous mutations promoted splicing changes primarily by SRE motifs and exonic synonymous and intronic mutations by RNA structure around the exon-intron junction. A combined model that accounted for both structure and SREs was the most accurate predictor of Exon 10 PSI, and most experimentally validated mutations altered RNA structure and SRE motif strength around the Exon 10-Intron 10 junction in the same direction (Figure 5D). The model further suggested that the overall region favored increased Exon 10 inclusion (Figure 6 A,B), which confirmed previous experimental findings that inclusion is Exon 10’s typical splicing mode (Q. S. Gao et al. 2000). This preference was proposed to be due to a weak 5’ splice site (Ian D’Souza and Schellenberg 2005), and, indeed, we found that almost all experimentally validated mutations strengthened the splice site to increase inclusion of Exon 10 (Figure 4A). However, interestingly, our model revealed that structural changes caused by the mutations resulted in a more structured exon-intron junction, which would imply decreased Exon 10 inclusion. However, SRE strength alterations overall skewed more towards increased Exon 10 inclusion, which suggest that SREs and the RBPs that bind them buffer the effects of RNA structure to maintain the 1:1 isoform ratio at this junction. Our work revealed that structure and splicing regulatory elements most often have opposite effects on splicing outcomes. However, disease variants were the exception to this rule and resulted in a synergistic effect on splicing outcome (Fig. 6E), leading to a greater disruption of splicing, and therefore increased pathogenicity. The combined model was finally validated by accurate prediction of the effects of six previously untested VUSs on Exon 10 splicing (Figure 6E). As was the case with the complete mutagenesis, there were few VUSs predicted to completely alter the ratio of isoforms to entirely 3R: only 5 VUSs had PSIs less than 0.25. However, our model accurately predicted the effect of the three 3R VUSs tested. Interestingly, the systematic computational mutagenesis revealed a hotspot of 3R mutations around 25-30 nucleotides downstream of the exon-intron junction (Figure 6A) and indeed the 3R VUSs experimentally validated were located in this region.

### Quantitative modeling of splicing regulation at exon-intron junctions

Predictive models can measure the contribution of individual factors to an outcome. Structure around the 5’ splice site and SRE motifs were excellent predictors of Exon 10 splicing in cells. The use of general SRE motifs enables this splicing framework to extend to other exon-intron junctions. By using a common dependent variable of Exon 10 PSI, we could use experimentally validated mutation data from disparate sources. Although our model provided an exact PSI prediction for each mutation, its principal utility was in predicting the direction in which the 3R:4R isoform ratio shifted from the wild type balance. On the basis of RNA-sequencing of brain tissue from healthy individuals, we find a range of Exon 10 PSIs between individuals and between tissues within an individual (Figure 1A). Even in individuals with progressive supranuclear palsy, a tauopathy in which *MAPT* variants are implicated, there is variability in Exon 10 PSIs between different brain tissues (Majounie et al. 2013). Ultimately, although it is likely that what is considered the correct ratio for normal brain function varies between tissues, our model provides a means to determine the baseline change of Exon 10 splicing simply based on sequence features. Many neurodegenerative diseases are caused by mutations around the *MAPT* Exon 10-Intron 10 junction, and there are no approved therapeutics that target this junction. Our work suggests that it is crucial to consider the larger structural context of the Exon 10-Intron 10 junction and the interplay between structure and SREs when considering the consequences of mutations on splicing regulation and the design of potential therapeutics to alter this ratio.

## Materials and methods

### *MAPT* Exon 10 PSIs for GTEx tissue types

Aligned BAM files of individual samples from the Genotype-Tissue Expression (GTEx) v8 project, for tissue types with *MAPT* TPM greater than 10, were accessed in the AnVIL/Terra environment (Kumar 2020a). Reads aligning to *MAPT* were extracted in Terra (Kumar 2020b) and downloaded. Exon 10 PSIs were quantified per BAM file with Outrigger (Song et al. 2017) using *MAPT* transcriptome reference from Ensembl GRCh38. Only samples with at least 10 reads mapping across the Exon 10-Intron 10 junction were considered. Median values for each tissue type were calculated and then visualized on the brain diagram with R package, CerebroViz (Bahl, Koomar, and Michaelson 2017). Source file for Figure 1 provides Exon 10 PSI values for the 2,962 samples. An ANOVA test was run in R to test significance in variation between individuals versus within an individual (for individuals with MAPT expression in more than 7 tissues) (Supplementary file 1).

### Culture of T47D and SH-SY5Y cells

Mammary gland carcinoma cells (T47D) were cultured in RPMI 1640 medium, supplemented with 10% Fetal Bovine Serum (FBS) and 0.2 units/mL of human insulin at 37°C and 5% CO_2_. Bone marrow neuroblastoma SH-SY5Y cells were cultured in 1:1 mixture of 1X Minimum Essential Medium (MEM) and 1X F12 medium, supplemented with 10% FBS at 37 °C and 5% CO_2_.

### In vivo DMS-MaP probing for *MAPT* RNA

Approximately 20 million T47D cells and 30 million SHSY-5Y cells were harvested by centrifugation and resuspended in bicine buffered medium (300 mM Bicine pH 8.3, 150 mM NaCl, 5 mM MgCl_2_) followed by treatment with DMS (1:10 ethanol diluted) for 5 min at 37°C. For the negative control (unmodified RNA), instead of DMS, an equivalent amount of ethanol was added to the same number of T47D and SH-SY5Y cells. After incubation, the reactions were neutralized by addition of 200 µl of 20% by volume β-mercaptoethanol. Total RNA was extracted by Trizol (ThermoFisher Scientific), treated with TURBODNase (ThermoFisher Scientific), purified using Purelink RNA mini kit (ThermoFisher Scientific) and quantified with NanoDrop™ spectrophotometer.

### DMS-MaP cDNA synthesis, library construction and sequencing for *MAPT* RNA

Purified RNA (9 μg) was reverse transcribed using Random Primer 9 (NEB) and SuperScript II reverse transcriptase under error prone conditions as described in Smola et al., 2015. The resultant cDNA was purified using G50 column (GE healthcare) and subjected to second strand synthesis (NEBNext Second Strand Synthesis Module). Supplementary file 4 lists PCR primers used for library generation. The cDNA was amplified by the NEB Q5 HotStart polymerase (NEB). Secondary PCR was performed to introduce TrueSeq barcodes (Smola et al. 2015). All samples were purified using the Ampure XP (Beckman Coulter) beads and quantification of the libraries was performed with Qubit dsDNA HS Assay kit (ThermoFisher Scientific). Final libraries were run on Agilent Bioanalyzer for quality check. TrueSeq libraries were then sequenced as necessary for their desired length as paired end 2×151 and 2×301 read multiplex runs on MiSeq platform (Illumina) for pre-cursor and mature MAPT isoforms respectively. Sequenced reads have been uploaded to the NCBI SRA database under BioProject ID PRJNA762079.

### In vivo DMS-MaP probing for *SSU ribosome*

For in vivo ribosomal structure data, we used approximately 10 million T47D cells in 10 cm plates for each condition. We removed the growth media, added 1.8 mL of bicine buffered growth medium (200 mM Bicine pH 8.3) followed by treatment at 37°C with 200 uL of DMS diluted in ethanol (1.25% final DMS) for 5 min. For the negative control (unmodified RNA), instead of DMS, an equivalent amount of ethanol was added to the same number of T47D cells. After incubation, all reactions were neutralized by addition of ice cold 20% by volume β-mercaptoethanol and kept on ice for 5 minutes. Total RNA was extracted by Trizol (ThermoFisher Scientific), chloroform and isoamyl alcohol using phase lock heavy tubes (5PRIME Phase Lock Gel). RNA was purified using Purelink RNA mini kit (ThermoFisher Scientific), treated with TURBODNase (ThermoFisher Scientific) and quantified with NanoDrop™ spectrophotometer.

### DMS-MaP cDNA synthesis, library construction and sequencing for *SSU ribosome*

Purified RNA (5 ug) was reverse transcribed using Random Primer 9 (NEB) and SuperScript II reverse transcriptase under error prone conditions as described Smola et al., 2015. The resultant cDNA was purified using G50 column (GE healthcare) and subjected to second strand synthesis (NEBNext Second Strand Synthesis Module). Standard Nextera DNA library protocol (Illumina) was used to fragment the cDNA and add sequencing barcodes. All samples were purified using Ampure XP (Beckman Coulter) beads and quantification of the libraries was performed with Qubit dsDNA HS Assay kit (ThermoFisher Scientific). Final libraries were run on Agilent Bioanalyzer for quality check. Libraries were sequenced as paired end 2×151 read multiplex runs on MiSeq platform (Illumina). Sequenced reads have been uploaded to the NCBI SRA database under BioProject ID PRJNA762079.

### DMS-MaP analysis

Sequenced reads were analyzed using the ShapeMapper pipeline(Busan and Weeks 2018), version (v2.1.4) which calculates the DMS reactivity of each nucleotide *i* as follows:

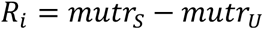

where *mutr_S_* is the mutation rate in the sample treated with DMS, *mutr_U_* is the mutation rate in the untreated control. DMS reactivities were normalized within a sample and per nucleotide type (A, C, U, G) using the normalization method described in Busan and Weeks, 2018. DMS probing data were collected for the Exon 9-Exon 11 and Exon 9-Exon 10-Exon 11 junctions using a single pair of primers listed in Supplementary file 4. The ShapeMapper pipeline ran for the two junctions in a single run with reference sequences for both junctions entered in one FASTA file. For the SSU, sequenced reads were first aligned to the SSU ribosome sequence using Bowtie2 parameters from Busan and Weeks, 2018. Using samtools, alignments with MAPQ score greater than 10 were kept, sorted, and converted back into FASTQ files after which the ShapeMapper pipeline was executed.

### Updating DMS parameters for RNAstructure using SSU ribosome data from T47D cells

To use DMS data to guide secondary structure prediction by the Rsample (Spasic et al. 2018) component of RNAstructure (Reuter and Mathews 2010), we calibrated the expected DMS reactivities per nucleotide. Using the SSU ribosome mapping data and the known secondary structure (Petrov et al. 2014), we determined histograms for DMS reactivities for unpaired nucleotides, nucleotides paired at helix ends, and nucleotides paired in base pairs stacked between two other base pairs. These DMS histograms can be invoked by Rsample with the “--DMS” command line switch as part of RNAstructure 6.4 or later. The histograms had long tails to relatively high reactivities. We empirically found that limiting reactivities in the histograms and in the input data to a reactivity of 5 (where higher values are set to 5) gave the best performance at improving SSU rRNA secondary structure. The “--max 5” parameter is used with Rsample to apply this limitation. Pre-release of Rsample code including in vivo DMS parameters is included as a zip file for review, and will be included in RNAstructure 6.4.

### Base-pairing probabilities for SSU

The partition function for the SSU was generated using Rsample, using either the sequence or using the sequence and the DMS reactivities. All possible *i*-*j* base pairing probabilities were summed for each nucleotide *i* to generate a base pairing probability per nucleotide *i*.

### ROC curves for predicting SSU base pairs

Using the known secondary structure of the SSU, we assigned a nucleotide as either 0 or 1 if it was paired or unpaired. DMS reactivities were used to predict whether a nucleotide was paired; the higher the DMS reactivity, the more likely a nucleotide is unpaired. Base pairing probabilities were subtracted from 1 to obtain the probability that a base was unpaired, with 0 implying base was paired and 1 implying that base was unpaired. ROC curves and AUC values were generated using the plotROC (Sachs 2017) R package.

### Arc plots

Arc plots were generated using Superfold (Siegfried et al. 2014) modified to process DMS reactivity data.

### Generating structural ensemble of Exon 10-Intron 10 *MAPT* junction

The partition function of the Exon 10-Intron 10 MAPT junction for wild type (WT) and mutations was calculated with DMS reactivities as restraints using Rsample (Spasic et al. 2018). The DMS reactivities, which were collected for the WT sequence, were also used for the mutations to restrain the structural space with the reactivity made NA at the nucleotide where the mutation occurred. The program stochastic (Reuter and Mathews 2010) was used to sample 1000 structures (in CT format) from the Boltzmann distribution wherein the likelihood a structure is sampled was proportional to the probability that it occurred in the distribution (Y. Ding and Lawrence 2003).

### t-SNE visualization of structural ensembles of WT, 3R and 4R mutations of Exon 10-Intron 10 *MAPT* junction

Structural ensembles were generated as described above for WT, 3R, and 4R mutations. For each sequence, the 1000 structures in CT format were converted to dot-bracket (db) format with ct2dot (Reuter and Mathews 2010), after which the db structure was transformed into the element format using rnaConvert in the Forgi package (Kerpedjiev, Höner Zu Siederdissen, and Hofacker 2015). In the element format, every base is represented by the subtype of RNA structure in which it is found: stem (s), hairpin (h), loop(m), 5’end(f), and 3’end(t). Hence, each db structure is a string of characters. These characters were digitized (f, t:0, s:1, h:2, m:3) to create a numerical matrix with 1000 rows and 234 columns, the length of the exon-intron junction.

Combining the matrices for the three sequences resulted in a 3000×234 matrix. This matrix was entered into the tSNE function from the scikit-learn python package (Pedregosa et al. 2011) and dimensionality was reduced to a 3000×2 matrix which was then plotted with ggplot2 (Wickham 2016) in R. The ΔG^‡^ of unfolding of the splice site was calculated for each of the 3000 structures as described below. Source file for Figure 3B lists t-SNE reduced data with corresponding free energies.

### Determining representative structures for clusters in t-SNE plot

The 3000×2 matrix, the result of t-SNE dimensionality reduction, was clustered using k-means clustering with the k-means function from the scikit-learn python package (Pedregosa et al. 2011). The value of k was set to 5 as determined visually. A custom python script was used to deduce the representative structure for each cluster by first calculating the most common RNA structure subtype at each position. The actual structure in the ensemble, most similar to the RNA structure with the most common subtypes at each position, was then determined and deemed to be the representative structure of that cluster.

### Visualizing density of structures in t-SNE plot

A custom python script was written. For the WT and, 3R, and 4R mutant sequences, a meshgrid was created for the three matrices using a 1000-point interpolation and NumPy (Harris et al. 2020) meshgrid function which returns two two-dimensional arrays that represent all the possible x-y coordinates for the three matrices. A gaussian kernel was next fit and evaluated for each 1000×2 matrix with SciPy gaussian_kde function (Virtanen et al. 2020) to smoothen over the meshgrid. Contour lines were generated for the smoothed data with Matplotlib contour function (Hunter 2007) and contourf was used to plot the data.

### Quantifying nucleotides around the 5’ splice site in cryo-EM structure

The Protein Databank (PDB) files for Pre-B (PDB ID: 6QX9), B (PDB ID: 5O9Z), Pre-B^act^ (PDB ID: 7ABF) and B^act^ (PDB ID: 5Z56) complexes were downloaded from the PDB website. A custom python script was used to extract pre-mRNA from each PDB file. The number of nucleotides were counted for mRNA found near the 5’ splice site. The result was visually confirmed by visualizing the PDB on PyMol.

### Calculating ΔG**^‡^** of unfolding of a region of interest

The ΔG^‡^ of a structure was calculated using the efn2 program in RNAstructure (Reuter and Mathews 2010). This represents the non-equilibrium unfolding energy of the region as the sequence is not allowed to refold after unfolding (Mustoe et al. 2018). The base pairs within a region of interest were removed using a custom python script. The ΔG^‡^ of the “unfolded” structure was next re-calculated with efn2. The ΔG^‡^ of unfolding of a region was the subtraction of the ΔG^‡^ of the original structure from the ΔG^‡^ of the unfolded structure. For example, for determining the ΔG^‡^ of unfolding of the splice site, we removed all base pairs within the last 3 nucleotides of the exon and the first 6 nucleotides of the intron.

### Calculating the change in strength of SRE motifs

#### Splice Site

Strength of the WT splice site was calculated with MaxEntScan (Yeo and Burge 2004). Strength was recalculated in the presence of splice site mutations either in the last 3 bases of Exon 10 or first 6 bases of Intron 10. WT strength was subtracted from the mutant strength: a 0 implied no change in splice site strength, positive values implied that a mutation made splice site stronger, resulting in increased inclusion of Exon 10, and negative values implied that a mutation made splice site weaker and decreased inclusion of Exon 10.

#### Enhancers and Silencers

Overrepresented hexamers in cell-based screens of general exonic and intronic splicing enhancers (ESEs, ISEs) and silencers (ESSs, ISSs) were obtained from Fairbrother et al., 2002, Wang et al., 2004, Wang, Ma et al., 2012 and Wang, Xiao et al., 2012. Position weight matrices (PWMs) of hexamers for each category were re-calculated as described in Fairbrother et al., 2002 and collated in Supplementary file 5. There were 8 clusters of ESE motifs, 7 of ESS motifs, 7 of ISE motifs, and 8 clusters of ISS motifs; each cluster had an associated PWM. For each PWM, a threshold strength was found by taking the 95^th^ percentile value of strength of all possible k-mers of PWM length. This threshold was used to determine whether there was a valid SRE motif at a particular position. The strength of the PWM motif was calculated across the exon-intron junction using a sliding window for both WT sequence and per mutation. The only windows that differed were around the location of the mutation, and only windows with strength above the threshold were considered. The WT strength was subtracted from the mutation strength for each window, and all windows were then summed to yield a Δstrength for every PWM per mutation. The average of the non-zero Δstrengths was calculated for ESE, ESS, ISE and ISS categories. The ESE and ISE Δstrengths were summed to obtain an enhancer strength, and the ESS and ISS Δstrengths were summed to obtain a silencer strength. Supplementary file 6 presents all SRE Δstrengths for the 47 mutations and 55 VUSs.

### Calculating the change in strength of RBP motifs

Position Frequency Matrices (PFMs) were available from Ray et al. 2013 for the following RBPs: SRSF1, SRSF2, SRSF7, SRSF9, SRSF10, PCBP2, RBM4 and SFPQ. PFMs were converted into PWMs by normalizing frequencies to 0.25 (Prior probability for nucleotide frequency) and calculating the log2 value. Overrepresented hexamers were available from Dominguez et al., 2018 for the following RBPs: SRSF11, SRSF4, SRSF5 and SRSF8. PFMs for those RBPs were calculated as described in Fairbrother et al., 2002. Δstrength in RBP motifs were calculated the same way as SRE motifs. The average of non-zero values of RBPs implicated in either the inclusion or exclusion of Exon 10 was computed separately. All RBP Δstrengths for the 47 mutations are found in Supplementary file 6.

### Models and bootstrapping

Exon 10 PSI was limited to values between 0 and 1 with 0 signifying that no transcripts had Exon 10 and 1 implying that all transcripts had Exon 10. Hence, standard linear regression was no longer appropriate and features were fit with a beta regression model to Exon 10 PSI. Regression parameters were determined using the betareg package (Cribari-Neto and Zeileis 2010) in R. Bootstrapping was performed by sampling without replacement 70% of the mutations to train and test the model and calculating the Pearson correlation coefficient (R^2^) between true values and predictions of the sample. This bootstrapping was executed 10 times resulting in a range of R^2^s, ensuring that no subset of mutations skewed model performance. Since there were only 4 mutations that maintained the wildtype 3R to 4R ratio in our training set, we added 3 additional variants of unknown significance (VUSs) from the Single Nucleotide Polymorphism database (dbSNP) which we experimentally verified preserved the wildtype splicing pattern (see Supplementary file 7 for gel). WT VUSs tested and added to the training set were assigned a PSI of 0.5 to indicate equivalence to the WT sequence. Eq. **1**, the structure ensemble model, uses four characteristics describing ***X***, the ΔG° of unfolding of the region of interest around the exon-intron junction for 1000 structures in the ensemble. Eq. **2**, the minimum free energy (MFE) model, uses just ***Y***, the ΔG° of unfolding of the exon-intron junction found within the spliceosome at the B^act^ stage for the single MFE structure. Eq. **3**, the splice site model, uses the difference in splice site strength between WT sequence and a mutation where *SS* represents splice site. Eq. **4**, the combined SRE model, uses the difference in SRE strength between WT sequence and a mutation where *SS* represents splice site, *E* represents enhancer, and *S* represents silencer. Eq. **5**, the RBP model, uses the difference in RBP motif strength between WT sequence and a mutation where *Ex* represents RBPs involved in the exclusion of Exon 10 and *In* represents RBPs involved in the inclusion of Exon 10. Eq. **6** is the interactive model between structure and SRE, and Eq.**7** is the additive model. *isNonSynonymous*, *isSynonymous* and *isIntronic* represent the category of mutation and is either 0 or 1. Supplementary file 6 summarizes the performance of the models and features utilized.

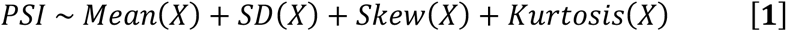

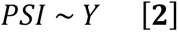

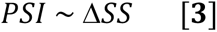

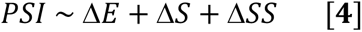

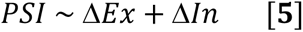

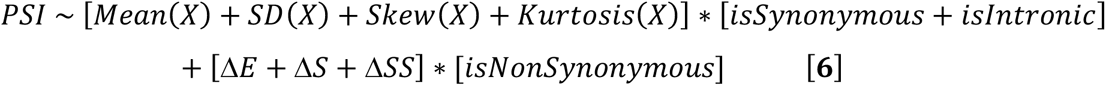

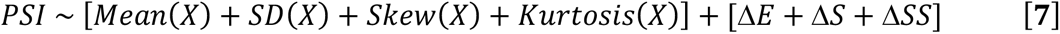

### Clustering changes in structural and SRE features

For each feature, non-zero values greater than the 95^th^ percentile value were set to the 95^th^ percentile or, if less than the 5^th^ percentile value, were set to the 5^th^ percentile for visualization, after which all values were normalized to the maximum absolute value. Silencer Δstrength and mean ΔG° of unfolding of exon-intron junction of ensemble were inverted to follow the visualization such that values closer to 1 would result in greater Exon 10 inclusion and values closer to 0 would result in lower Exon 10 inclusion. Features were then assigned values 0 or 1 depending on whether the feature changed at all in the presence of the mutation. These digitized features were first clustered by hierarchal clustering resulting in 6 clusters. Each individual cluster was clustered again by hierarchal clustering but using the normalized feature values instead of 0s and 1s.

### Splicing Assays

HEK-293 cells (ATCC CRL-1573) were grown at 37°C in 5% CO_2_ in Dulbecco’s Modified Eagle Medium (Gibco) supplemented with 10% FBS (Omega Scientific) and 0.5% Penicillin Streptomycin (Gibco). The wild type splicing reporter plasmid was generously provided by the Roca lab and is described in Tan et al., 2019. Single-nucleotide point mutations were generated using a Q5 site-directed mutagenesis kit (NEB) and confirmed by Sanger sequencing, or custom ordered directly from GenScript. Reporter plasmids (2 μg) were transfected into HEK-293 cells in 6-well plates at ∼60-90% confluency using Lipofectamine 3000 (ThermoFisher Scientific). Cells were harvested after 1 day by aspirating the media and resuspending the cells in 1 mL Trizol reagent (ThermoFisher Scientific). RNA was isolated using the PureLink RNA Isolation Kit (ThermoFisher Scientific) with on-column DNase treatment, following manufacturer’s instructions. RNA (1 μg) was reverse transcribed to cDNA using Superscript VILO reverse transcriptase (ThermoFisher Scientific). Reverse transcriptions were performed by annealing (25°C 10 minutes), extension (50°C 10 minutes), and inactivation (85°C 10 min) steps. Heat-inactivated controls were prepared by heating the reaction without RNA at 85°C for 10 minutes prior to adding RNA, then following the described reaction conditions. The cDNA was PCR amplified with NEB Q5 HotStart polymerase (NEB) using splicing assay primers from IDT (AGACCCAAGCTGGCTAGCGTT forward, GAGGCTGATCAGCGGGTTTAAAC reverse) with 25 cycles. PCR product was purified and concentrated using the PureLink PCR micro clean up kit (ThermoFisher Scientific), following manufacturer’s instructions. Splicing products were visualized by loading ∼200 ng of DNA on a 2% agarose gel in 1X tris-acetate EDTA (TAE) buffer and staining with ethidium bromide. Gel images were quantified with ImageJ.

**Supplementary files, figure source files, SNRNASMs and code are available at GitHub repository: https://git.io/JuSW8**

## Acknowledgements

This work was supported by the US National Institutes of Health R01 HL111527 and R35 GM 140844 to A.L. and R01 GM076485 to D.M. The authors wish to thank the Roca Lab for providing wildtype splicing reporter plasmids, Dr. Zefeng Wang for intronic splicing enhancer and silencer motifs, and Drs. Peter Castaldi, John Platig and Kevin Weeks for insightful discussions.

## Supplementary Figures

**Figure 1-figure supplement 1:**
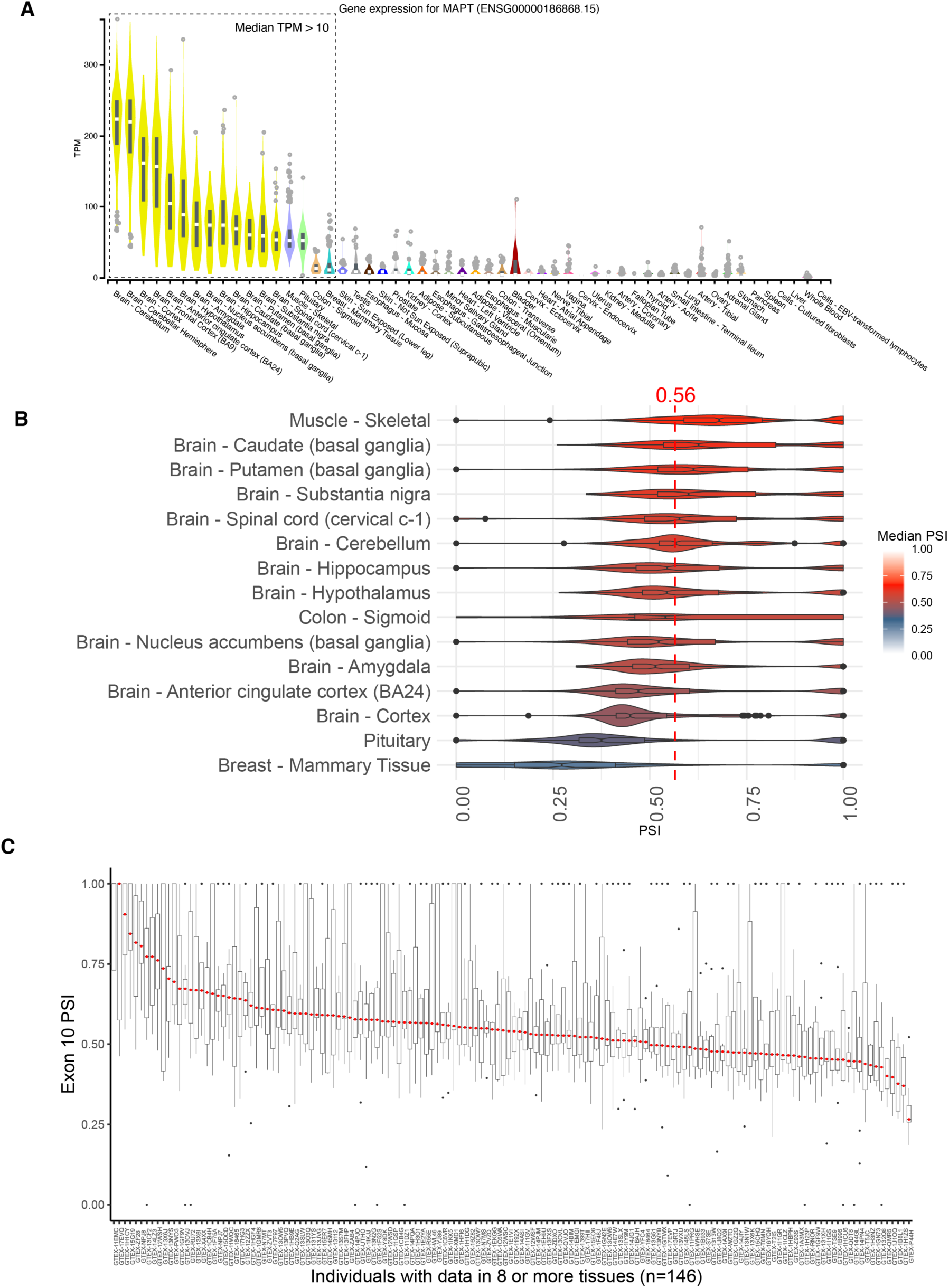
Distribution of Exon 10 PSIs calculated for RNA-seq data from GTEx database. A) Distribution of TPM values of *MAPT* gene expression for tissues in the GTEx database sorted by median TPM. Dotted box indicates tissues with median TPM greater than 10. MAPT is highly expressed in the brain, and there is little expression in other tissues. Figure was downloaded from the GTEx website. B) Distribution of Exon 10 PSI for 12 central nervous system, muscle-skeletal, colon-sigmoid, and breast-mammary tissue types. Percent Spliced In (PSI) of Exon 10 was calculated from RNA-seq data for 2,962 tissue samples among 15 tissue types collected from 818 individuals in GTEx v8 database. The violin plot for each tissue type and the corresponding region on the brain diagram is colored by the median PSI for all samples of that type. The median PSI of 0.56 for all tissue samples is indicated by the red dotted line. C) Distribution of Exon 10 PSI for tissues per individual. Only individuals with *MAPT* expression data in 8 or more tissues were plotted. Median PSI for each individual is labelled by red dot on box plot.

**Figure 1-figure supplement 2:**
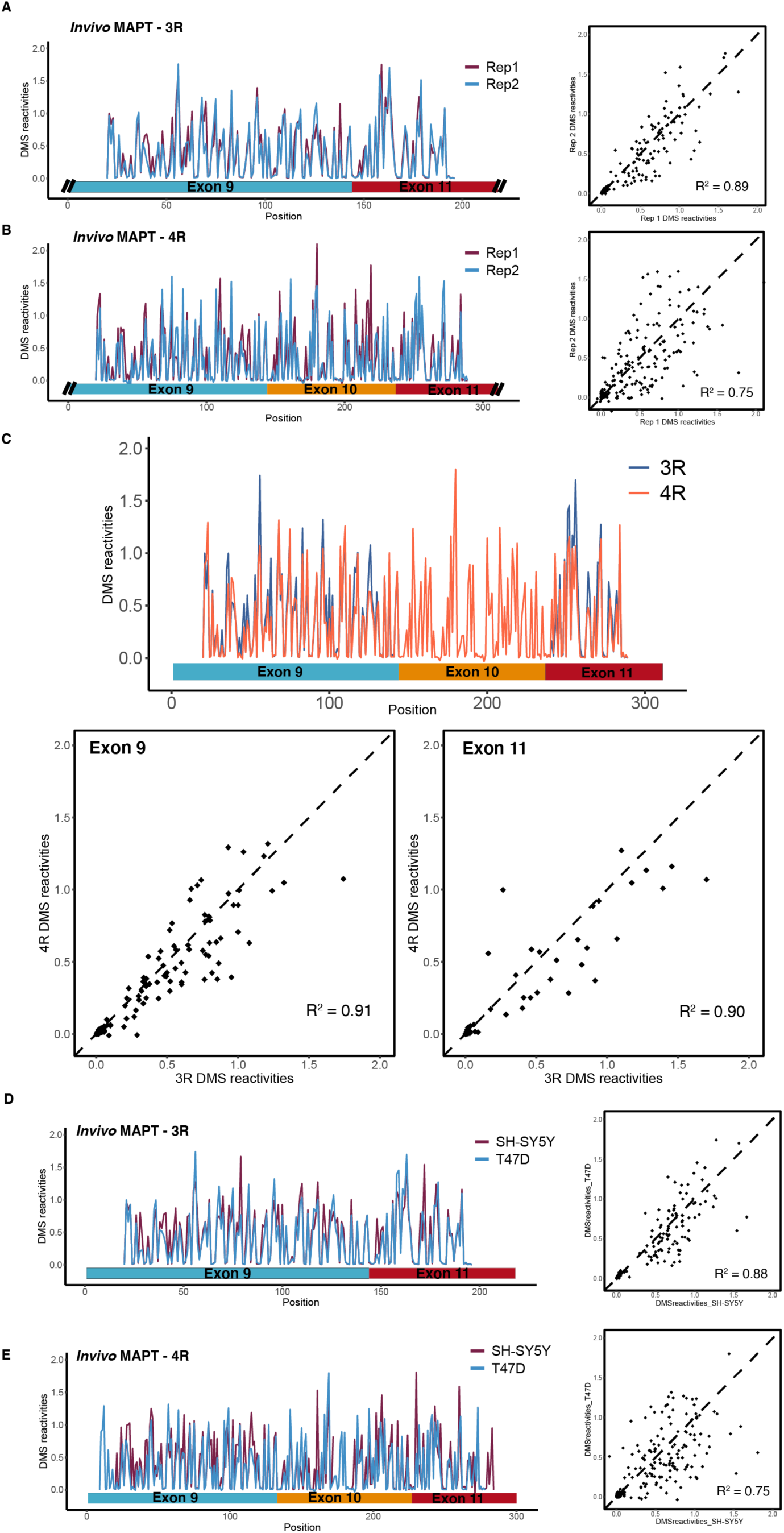
DMS structure probing data for mature *MAPT* 3R and 4R isoforms. A) DMS reactivity data from T47D cells for two biological replicates for Exon 9-Exon 11 junction (3R isoform). Structure probing data for junctions of interest were obtained using primers (Supplementary file 4) following RT of extracted RNA. DMS reactivity is plotted for each nucleotide across spliced junctions for both replicates overlaid in plot on the left. For scatter plot on the right, DMS reactivity for Rep 1 vs Rep 2 is plotted per nucleotide with Pearson’s correlation coefficient displayed. B) DMS reactivity data from T47D cells for two biological replicates for Exon 9-Exon 10-Exon 11 junction (4R isoform). C) Comparison of DMS reactivity data for 3R vs 4R isoforms. Replicates 1 and 2 were pooled for each isoform. Top plot shows DMS reactivity plotted for each nucleotide with isoforms overlaid. No data were collected for Exon 10 for the 3R isoform because Exon 10 is spliced out. Bottom left scatter plot shows DMS reactivities for Exon 9 in the 3R vs 4R context, whereas bottom right scatter plot shows DMS reactivities for Exon 11 in the 3R vs 4R context. Pearson’s correlation coefficient is shown for each comparison. D) DMS reactivity data from T47D and SH-SY5Y cells for Exon 9-Exon 11 junction (3R isoform). Replicates from T47D cells were pooled. DMS reactivity is plotted for each nucleotide across spliced junctions for both replicates overlaid in plot on the left. For scatter plot on the right, DMS reactivity for T47D vs SH-SY5Y is plotted per nucleotide with Pearson’s correlation coefficient displayed. E) DMS reactivity data from T47D and SH-SY5Y cells for Exon 9-Exon 10-Exon 11 junction (4R isoform).

**Figure 1-figure supplement 3:**
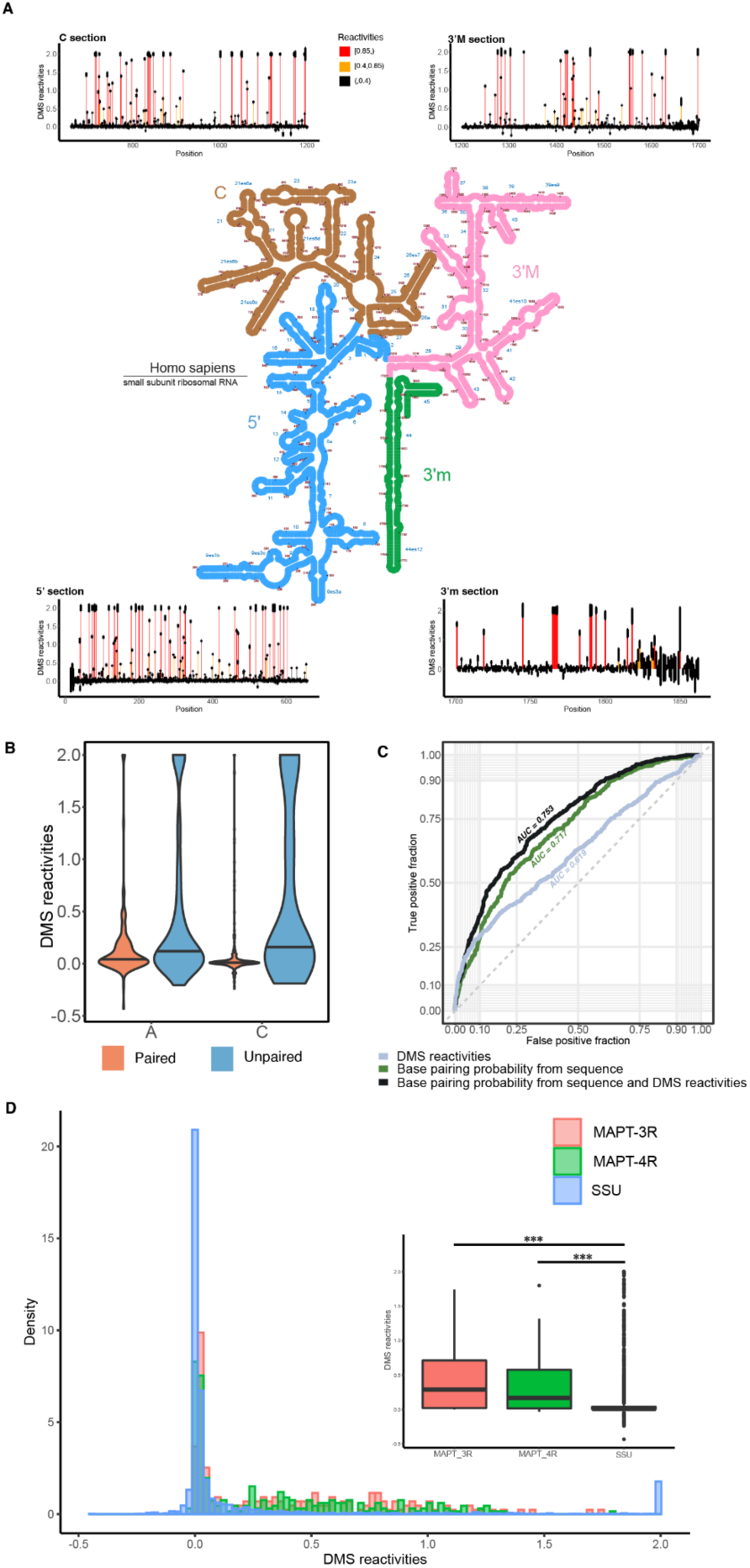
DMS structure probing data for human small subunit ribosome RNA (SSU) collected from T47D cells. A) T47D cells were treated with DMS. Data for SSU were extracted by aligning reverse transcribed RNA-seq data against the SSU sequence, after which reactivities were calculated. DMS reactivities are plotted for each of the four sub-domains of the SSU. Each value is shown with its standard error and colored by reactivity based on color scale. Reactivities were cut off at 2. High DMS reactivities correspond to unstructured regions, whereas low DMS reactivities correspond to structured regions. The secondary structure of the SSU was downloaded from Loren William’s lab Ribosome Gallery website (http://apollo.chemistry.gatech.edu/RibosomeGallery/eukarya/H%20sapiens/SSU/ind ex.html). B) Violin plots showing distribution of DMS reactivities for adenines and cytosines partitioned by paired versus unpaired nucleotides. Pairing status of nucleotides was determined from the known secondary structure of the SSU. Median DMS reactivity is indicated by thick horizontal black line on violin plot. C) ROC curves for predicting whether a nucleotide in the SSU is paired. Three different parameters were used for each of the three curves: DMS reactivities, base pairing probabilities predicted from SSU sequence, and base pairing probabilities for SSU sequence that were guided by DMS reactivities. The area under the curve (AUC) for each curve was calculated with AUCs closer to 1 corresponding to higher accuracy of predictions. Dotted line indicates AUC of 0.5 which corresponds to a model making random predictions. D) Comparison of distribution of DMS reactivities between SSU, *MAPT* 3R and 4R isoforms. Larger plot shows a density histogram of the DMS reactivities for each RNA. Inset boxplots display distribution of DMS reactivities. Level of significance: ***p-value < 10^-6^

**Figure 2-figure supplement 1:**
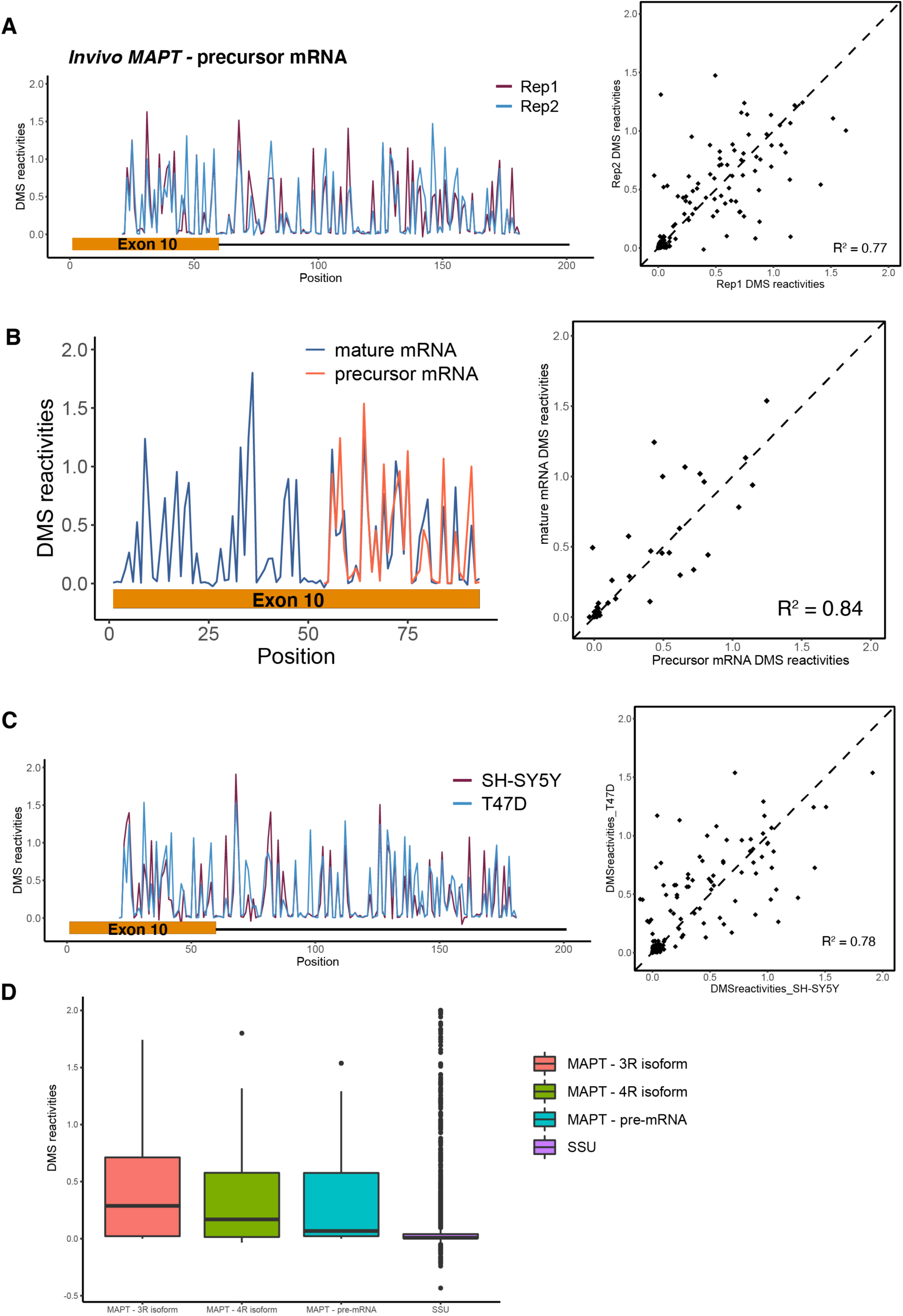
DMS structure probing data for precursor *MAPT* Exon 10-Intron 10 junction. A) DMS reactivity data from T47D cells for two biological replicates for Exon 10-Intron 10 junction. Structure probing data for junctions of interest were obtained using primers following RT of extracted RNA. DMS reactivity is plotted for each nucleotide across spliced junctions for both replicates overlaid in plot on the left. For scatter plot on the right, DMS reactivity for Rep 1 vs Rep 2 is plotted per nucleotide with Pearson’s correlation coefficient displayed. B) DMS reactivity data comparing Exon 10 in precursor vs mature transcript. Replicates 1 and 2 were pooled for each transcript. Right plot shows DMS reactivity plotted for each nucleotide with mature and precursor RNAs overlaid. DMS data for all of Exon 10 could not be collected for the precursor RNA due to the position of primers chosen for amplification. Scatter plot on the left shows DMS reactivities for Exon 10 in the precursor vs mature mRNA context with Pearson’s correlation coefficient shown for the comparison. C) DMS reactivity data from T47D and SH-SY5Y cells for Exon 10-Intron 10 junction. Replicates from T47D cells were pooled. DMS reactivities are plotted for each nucleotide across exon-intron junctions for both cell types overlaid in plot on the left. For scatter plot on the right, DMS reactivity for T47D vs SH-SY5Y is plotted per nucleotide with Pearson’s correlation coefficient displayed. D) Boxplots of distribution of DMS reactivities between SSU, *MAPT* 3R isoform, 4R isoform and pre-cursor mRNA.

**Figure 2-figure supplement 2:**
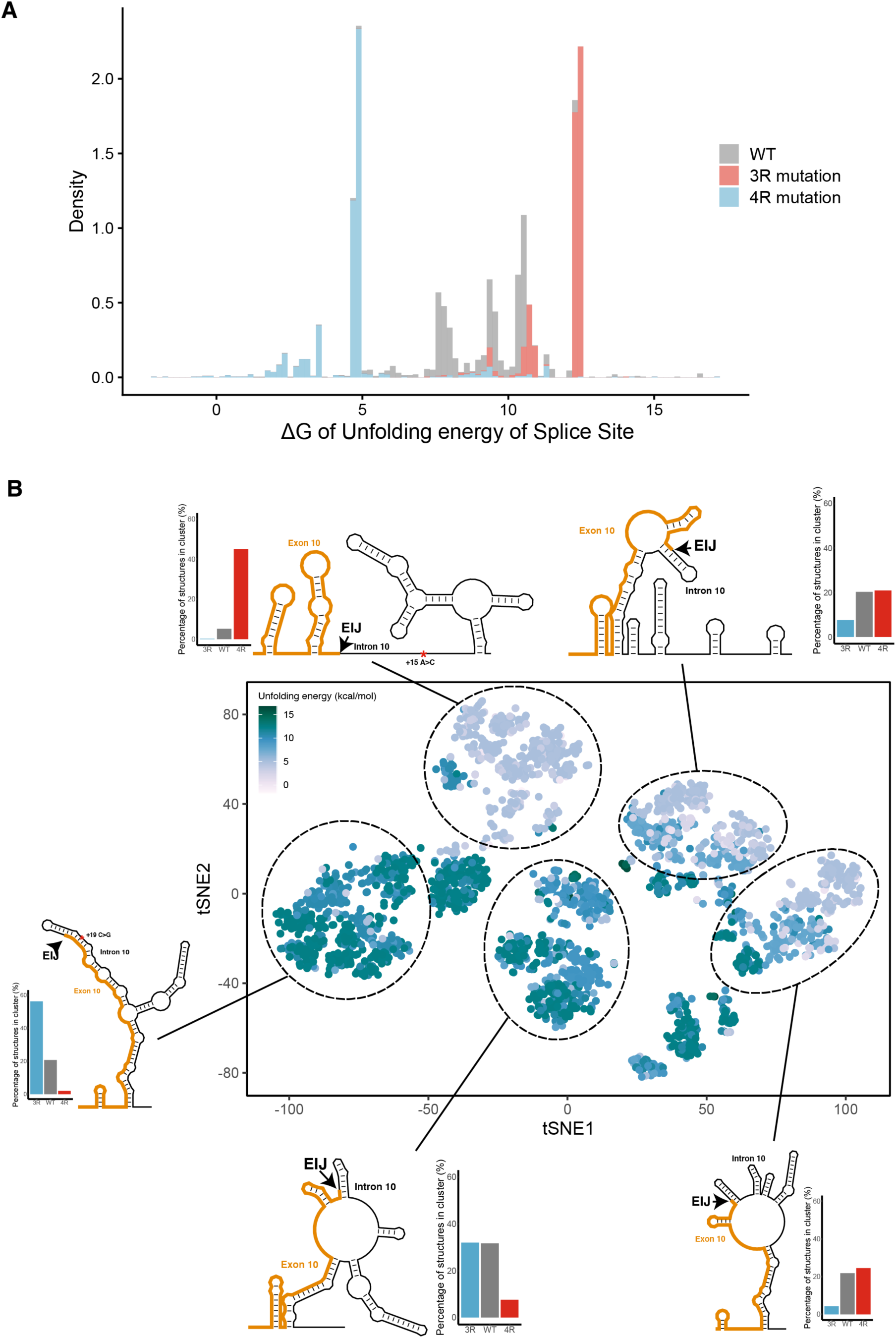
The 3R and 4R mutations shift the WT structural ensemble towards less and more accessible exon-intron junctions, respectively. A) Density histogram showing the distribution of unfolding free energies of the splice site (defined as last 3 nucleotides of exon, first 6 nucleotides of intron) for all structures in the ensemble for WT, 3R and 4R mutated sequence. Distributions for each sequence are colored according to the legend. B) t-SNE Visualization of structural ensemble of wild type (WT) and, 3R (+19C>G) and 4R (+15A>C) mutations. Structures were predicted using Boltzmann suboptimal sampling and guided by DMS reactivity data (in Figure 2A). Data were visualized using t-Distributed Stochastic Neighbor Embedding (t-SNE). Shown are 3000 structures corresponding to 1000 structures per category. Each dot represents a single structure and was colored by calculated unfolding free energy of splice site at exon-intron junction (3 exonic bases, 6 intronic bases). Data were clustered by k-means clustering and representative structures for five clusters are shown. Bar plots next to the representative structure show the proportion of the cluster in WT, 3R and 4R. The exon-intron junction is marked by EIJ on each structure. Position of 3R and 4R mutations are marked by a red asterisk on their respective representative structures. There are two additional representative structures shared by WT and 4R sequence which have similar structural contexts around the EIJ as the representative WT structure in Figure 2B.

**Figure 3-figure supplement 1:**
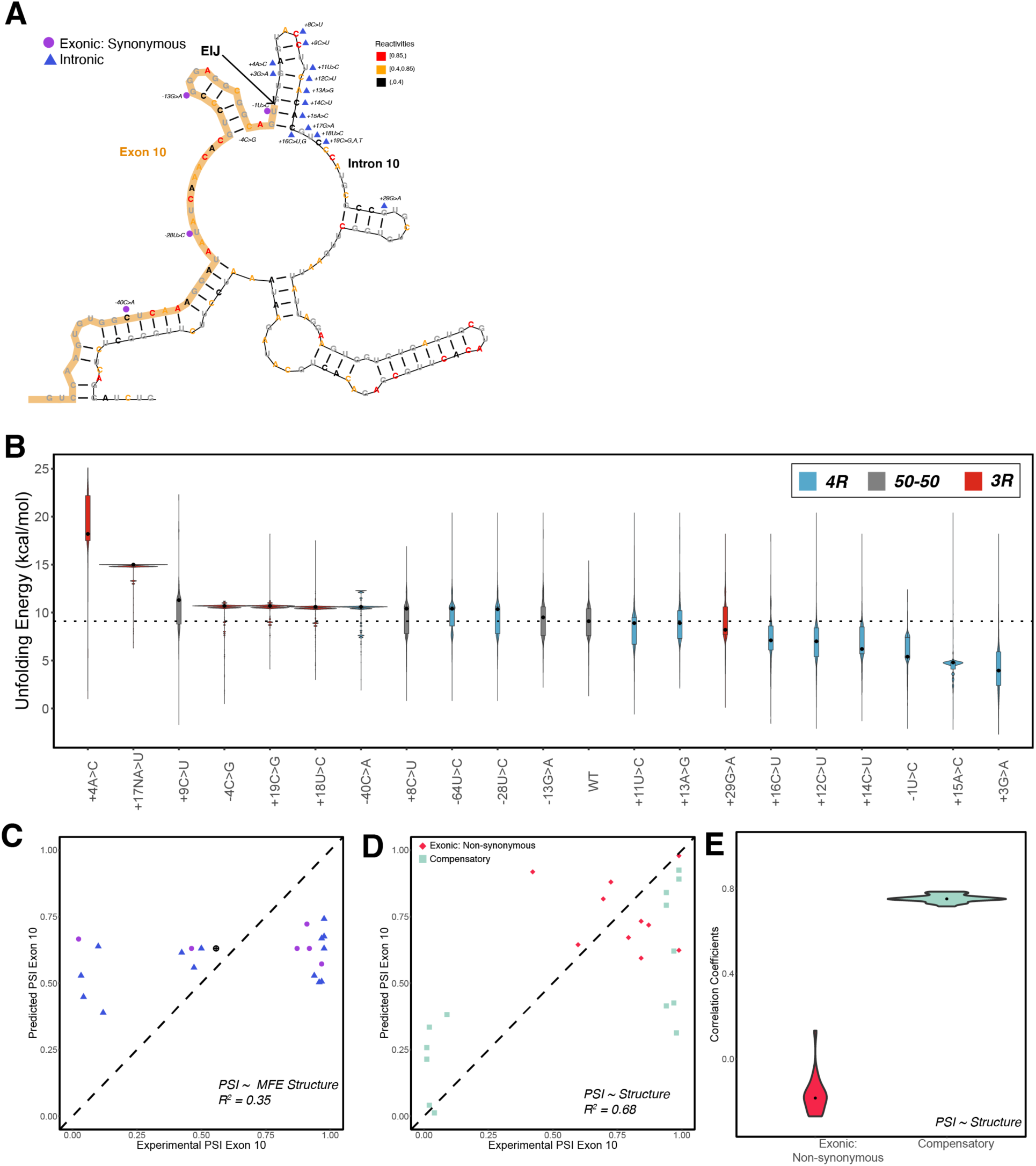
Structure is a poor predictor of Exon 10 PSI for exonic non-synonymous mutations. A) Positions of intronic and synonymous experimentally validated mutations, used in training the structure model, are shown on the wild type representative structure from Figure 2B. Each nucleotide is colored by its corresponding reactivity value based on the color scale. B) Violin plots showing the distribution of unfolding free energy of the exon-intron pre-mRNA in the spliceosome B^act^ complex for the 1000 structures in the ensembles of the 22 intronic and synonymous mutations. Each violin plot is colored by whether the mutation promotes the 3R or 4R isoform ratio or the ratio remains 50:50. C) Exon 10 PSIs of 22 mutations predicted using unfolding free energy of the exon-intron pre-mRNA in B^act^ complex of the spliceosome for the single minimum free energy (MFE) structure and plotted against experimental PSIs measured in splicing assays. Exon 10 PSIs predicted using Eq. 2. Each point on the scatterplot represents a mutation and is colored by mutation category. Dotted diagonal line is the x=y line, and the closer the points are to the diagonal, the more accurate the prediction. Pearson correlation coefficient (R^2^) of experimental to predicted PSIs was calculated. D) Exon 10 PSIs of non-synonymous and compensatory mutations predicted using the unfolding free energy of pre-mRNA within the spliceosome B^act^ stage plotted against corresponding experimental PSIs measured in splicing assays. Exon 10 PSIs were predicted using Eq. 1. E) Violin plots show R^2^s calculated for each mutation category by training and testing on subsets of all mutations by non-parametric bootstrapping; Non-synonymous (n=10), Compensatory (n=14).

**Figure 4-figure supplement 1:**
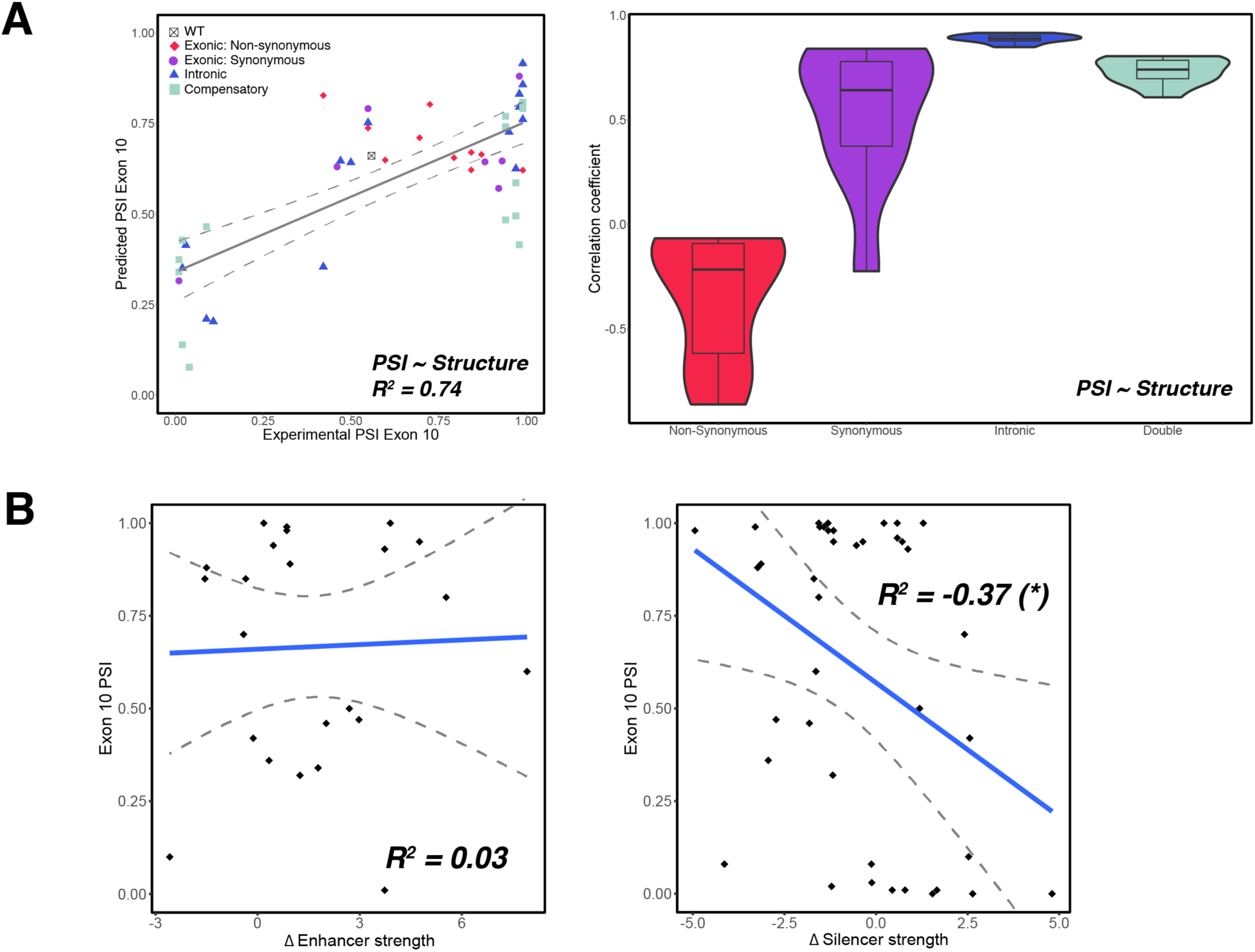
RBP binding motif strength is a poor predictor of Exon 10 PSI for all mutations. A) Exon 10 PSIs of 47 mutations predicted from structural change and plotted against experimental PSIs measured in splicing assays. Exon 10 PSIs predicted using Eq. 1. Each point on the scatterplot represents a mutation and is colored by mutation category. Grey line represents the best fit with dotted lines indicating the 95% confidence interval. Pearson correlation coefficient (R^2^) of experimental to predicted PSIs. Violin plot shows R^2^s calculated for each category by training and testing on subsets of all mutations by non-parametric bootstrapping; Exonic non-synonymous (n=11), Exonic synonymous (n=7), Intronic (n=15), Compensatory (n=14), Wildtype (n=1). B) Scatter plot of change in enhancer or silencer strength versus Exon 10 PSI. Each point represents a mutation. Blue line represents the line of best fit with dotted lines indicating the 95% confidence interval. Pearson correlation coefficient (R^2^) is shown. The negative correlation between silencer strength and Exon 10 PSI is statistically significant with a p-value of 0.004.

**Figure 4-figure supplement 2:**
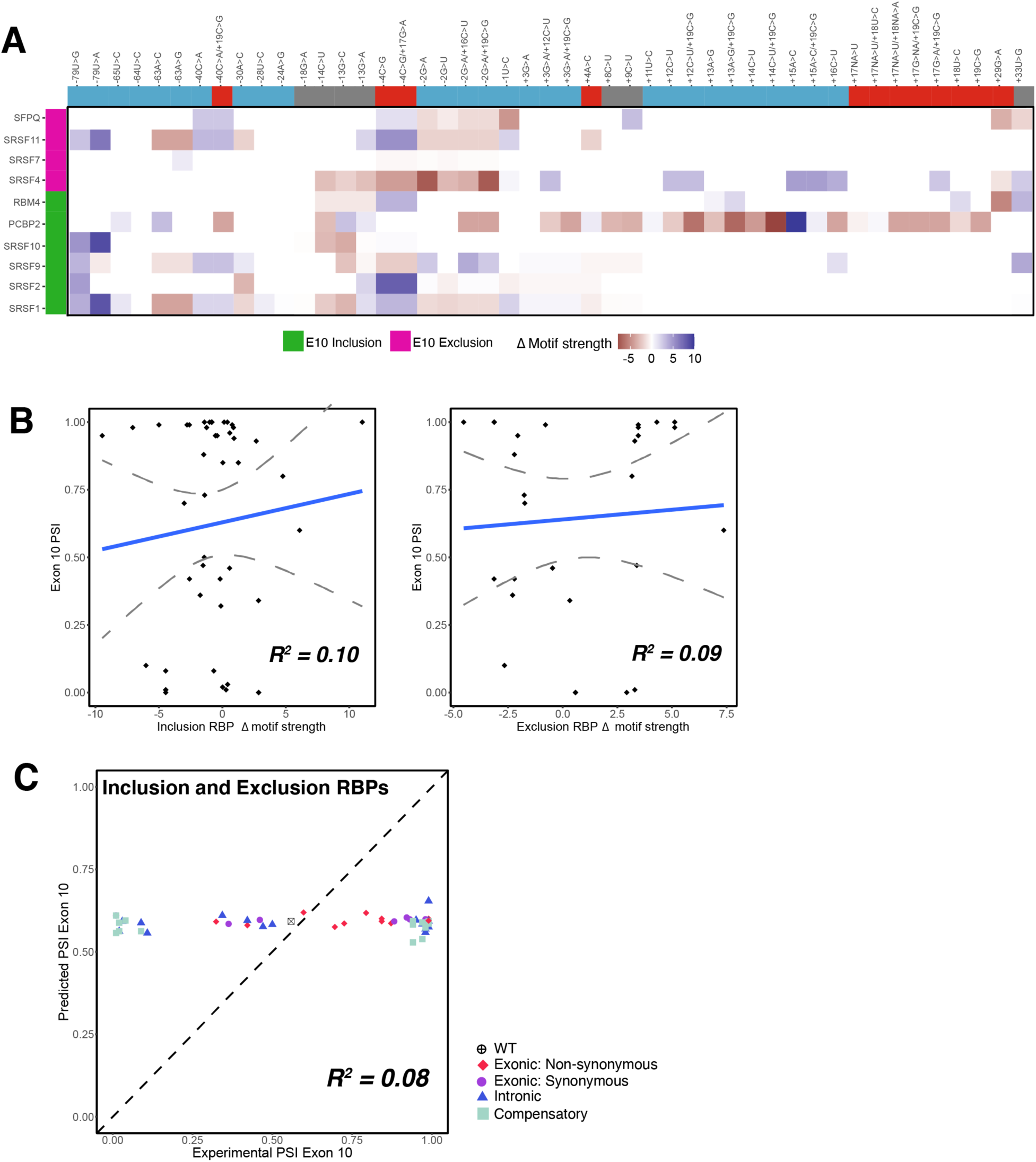
RBP binding motif strength is a poor predictor of Exon 10 PSI for all mutations. A) Heatmap of relative RBP binding motif strengths compared to wild type for 44 mutations. A value of 0 indicates that the mutation does not change RBP binding motif strength, a positive value indicates increase in RBP binding motif strength, and a negative value indicates weaker strength. RBPs implicated in the regulation of Exon 10 splicing were collected from Qian & Liu, 2014 and the RBP binding motifs were from Dominguez et al., 2018 and Ray et al., 2013. RBPs on the left, implicated in the splicing inclusion of MAPT Exon 10, are highlighted in pink, and RBPs involved in the exclusion of Exon 10 are highlighted in green. Mutations are marked based on whether they promote the 3R or 4R isoform ratio or the ratio remains 50:50. B) Scatter plot displaying change in RBP motif strength versus Exon 10 PSI, categorized based on whether the RBP is implicated in exclusion or inclusion of Exon 10. Neither correlation coefficient is statistically significant. C) Exon 10 PSIs of 44 mutations and wild type predicted using change in RBP motif strength and plotted against experimental PSIs measured in splicing assays. Exon 10 PSIs predicted using Eq. 5. Each point represents a mutation and is colored by category of mutation. Dotted diagonal line is the x=y line, and the closer the points are to the diagonal, the more accurate the prediction. Pearson correlation coefficient (R^2^) of experimental to predicted PSIs was calculated.

**Figure 5-figure supplement 1:**
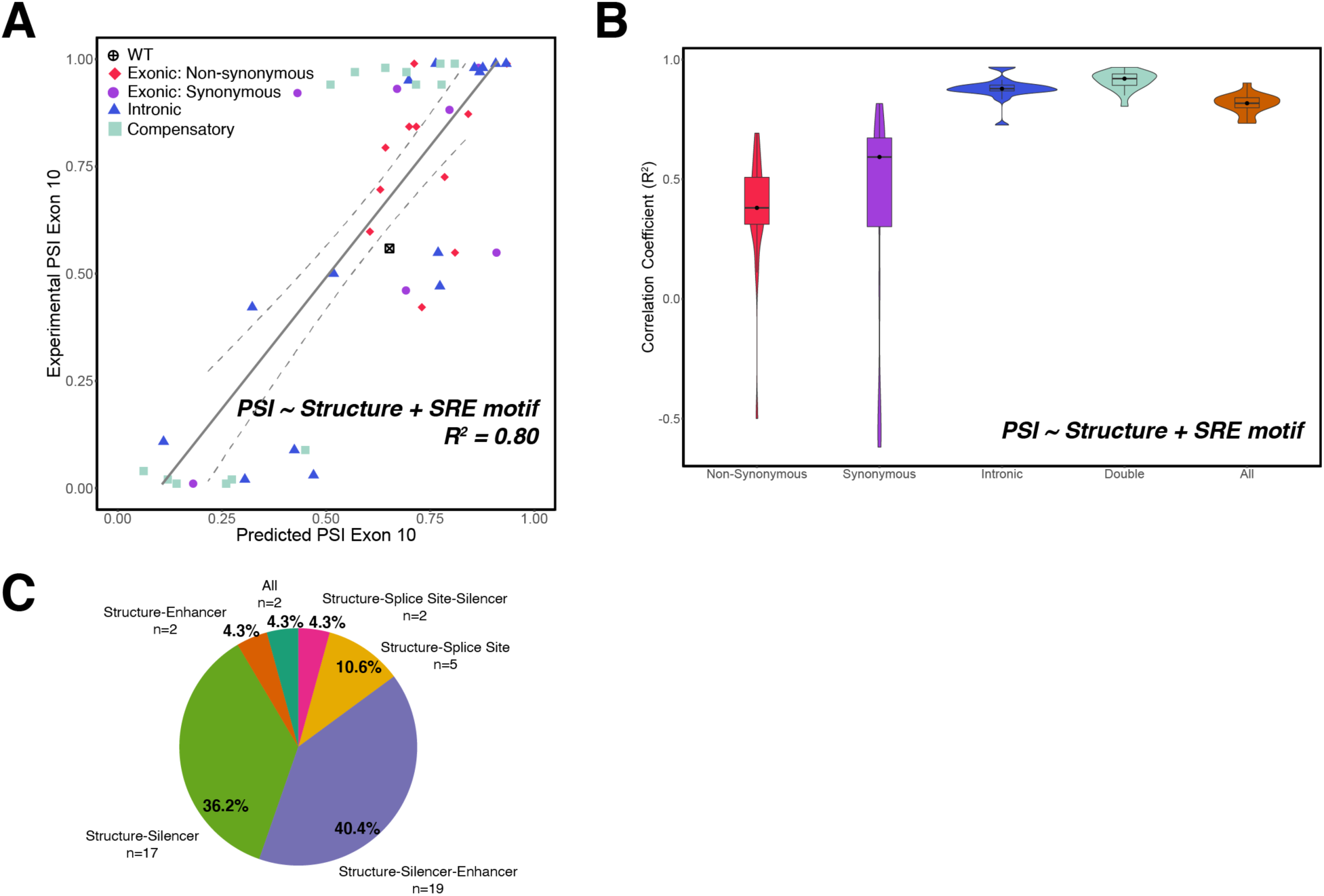
Additive model of structure and SRE has poorer predictive performance compared with an interactive model specifically for synonymous and non-synonymous mutations. A) Exon 10 PSIs of 47 mutations and wild type predicted using addition between structure and SRE strength and fit to experimental PSIs measured in splicing assays. Exon 10 PSIs are predicted using Eq. 7. Each point on scatterplot represents a mutation and is colored by category of mutation. Grey line represents the best fit with dotted lines indicating the 95% confidence interval. Pearson correlation coefficient (R^2^) of experimental to predicted PSIs was calculated. B) Violin plots showing correlation coefficients for each mutation category for structure and SRE additive model. R^2^s calculated for each mutation category by training and testing on subsets of all mutations by non-parametric bootstrapping 10 times. C) Pie chart of number and proportion of experimentally validated mutations in each cluster for heatmap in Fig 5B. Color of segment of pie chart matches up to the color of dendrogram branch in Fig 5B.

**Figure 6-figure supplement 1:**
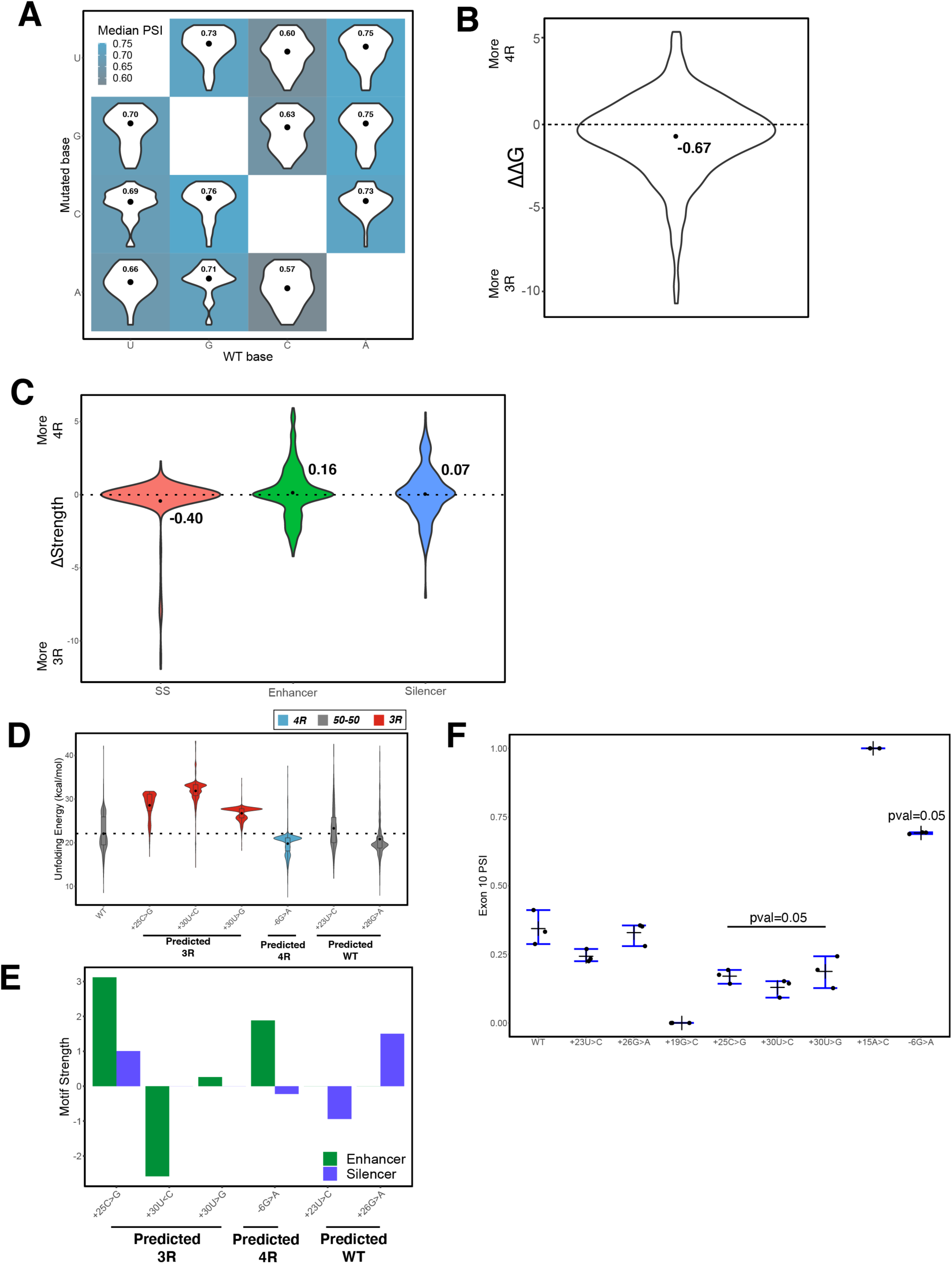
Complete mutagenesis of 100-nucleotide window spanning Exon 10-Intron 10 junction. A) Heatmap of mean predicted Exon 10 PSIs grouped by wild type and mutant nucleotide. Mutations were grouped by wild type and mutant nucleotide, and mean predicted PSIs were calculated by group and colored according to color scale. Violin plots of the distribution of PSI per group are shown in tile corresponding to group. On each tile, mean PSI is indicated by dot and labeled within violin plot. B) Violin plot of the distribution of normalized change in unfolding free energy of the exon-intron pre-mRNA in the spliceosome B^act^ complex from WT for all mutations around a 100-nucleotide window of exon-intron junction. Mean of -0.67 is indicated by dot. Dotted line represents the 0 value where there is no difference between WT and mutant unfolding free energy. Positive values imply region becomes less structured and has increased inclusion of Exon 10 (4R isoform); negative values are interpreted as more structured and decreased inclusion of Exon 10 (3R isoform). C) Violin plots showing the distribution of normalized change in splice site, enhancer, and silencer strength compared with WT for all mutations spanning a 100-nucleotide window of exon-intron junction. Mean is indicated by large black dots on violin plots. Dotted lines represent the 0 value where there is no difference from WT strength for mutation. Positive values suggest increased inclusion of Exon 10 (4R isoform), whereas negative values are interpreted as decreased inclusion of Exon 10 (3R isoform). D) Violin plot shows the distribution of unfolding free energy of the exon-intron pre-mRNA in the spliceosome B^act^ complex for the 1000 structures in the ensembles of wild type and the 6 VUSs experimentally tested. Each violin plot is colored by whether the mutation promotes the 3R or 4R isoform ratio or the ratio remains 50:50. The dotted line indicates the median unfolding free energy of the WT ensemble. E) Bar plots display the change in enhancer and silencer strength of the 6 VUSs compared with WT. F) Quantification of Exon 10 PSI of three replicates for splicing assay gels for 6 VUSs. One tailed Wilcoxon Rank Sum test was used to calculate significance of Exon 10 PSI of VUS of interest compared to WT.

## Supplementary Files

**Supplementary file 1**: ANOVA table for between individuals and within individuals Exon 10 PSI comparison

**Supplementary file 2**: Details on 47 experimentally tested *MAPT* mutations used in training model

**Supplementary file 3**: Details on 55 variants of unknown significance (VUSs) in *MAPT* from dbSNP

**Supplementary file 4**: Primers used for amplification of exon-exon or exon-intron junctions

**Supplementary file 5**: Re-calculated Position Weight Matrices for ESEs, ESSs, ISEs, ISSs

**Supplementary file 6**: Details on beta regression model results and features used for each training and test set

**Supplementary file 7**: Gel of RT-PCR data for splicing assay for new WT VUSs

